# CYCA3;4 Is a Post-Prophase Target of the APC/C^CCS52A2^ E3-Ligase Controlling Formative Cell Divisions in Arabidopsis

**DOI:** 10.1101/2020.03.17.995506

**Authors:** Alex Willems, Jefri Heyman, Thomas Eekhout, Ignacio Achon, Jose Antonio Pedroza-Garcia, Tingting Zhu, Lei Li, Ilse Vercauteren, Hilde Van den Daele, Brigitte van de Cotte, Ive De Smet, Lieven De Veylder

## Abstract

The Anaphase Promoting Complex/Cyclosome (APC/C) controls unidirectional progression through the cell cycle by marking key cell cycle proteins for proteasomal turnover. Its activity is temporally regulated by the docking of different activating subunits, known in plants as CDC20 and CCS52. Despite the importance of the APC/C during cell proliferation, the number of identified targets in the plant cell cycle is limited. Here, we used the growth and meristem phenotypes of Arabidopsis CCS52A2-deficient plants in a suppressor mutagenesis screen to identify APC/C^CCS52A2^ substrates or regulators, resulting in the identification of a mutant cyclin *CYCA3;4* allele. CYCA3;4 deficiency partially rescues the early *ccs52a2-1* phenotypes, whereas increased CYCA3;4 levels enhances them. Furthermore, whereas CYCA3;4 proteins are promptly broken down after prophase in wild-type plants, they remain present in later stages of mitosis in *ccs52a2-1* mutant plants, marking them as APC/C^CCS52A2^ substrates. Strikingly, *CYCA3;4* overexpression results in aberrant root meristem and stomatal divisions, mimicking phenotypes of plants with reduced RBR1 activity. Correspondingly, RBR1 hyperphosphorylation was observed in CYCA3;4-overproducing plants. Our data thus demonstrate that an inability to timely destroy CYCA3;4 attributes to disorganized formative divisions, likely in part caused by the inactivation of RBR1.

**ONE-SENTENCE SUMMARY:** Timely post-prophase breakdown of the Arabidopsis cyclin CYCA3;4 by the Anaphase Promoting Complex/Cyclosome is essential for meristem organization and development.

## INTRODUCTION

Cell division represents an essential biological process, not only allowing the transfer of genetic information from one generation to the next, but also permitting multicellular organisms to grow and develop. The latter implies the control of cell proliferation in such a way that a building plan can be carried out. When a new cell arises through cell proliferation from the stem cells, it frequently undergoes a number of cell divisions that are eventually followed by the execution of a cell cycle exit program. Both the proliferative activity of the stem cells and the timing of cell cycle exit need to be strictly regulated, as perturbations in either impair growth (De Veylder et al., 2007; Polyn et al., 2015; Shimotohno and Scheres, 2019). One of the key players that controls both events is the Anaphase Promoting Complex/Cyclosome (APC/C) (see Heyman and De Veylder (2012) for an extensive review on the plant APC/C). The APC/C is a conserved E3 ubiquitin ligase that provides unidirectional transit through the cell cycle by targeting key cell cycle proteins for degradation by the 26S proteasome (Marrocco et al., 2010). The plant APC/C consists of at least 11 core subunits, of which most are coded for by single-copy genes that are essential for plant viability (Page and Hieter, 1999; Capron et al., 2003; Van Leene et al., 2010; Heyman and De Veylder, 2012). Its structural backbone consists of the tetratricopeptide repeat (TPR) interaction domain-containing proteins APC6, APC7, APC8 and APC3 (the latter being present in two copies in *Arabidopsis:* APC3a/CDC27 and APC3b/HOBBIT) and is completed by APC1, APC4 and APC5. Together, they correctly position the catalytic subunits APC2 and APC11, which perform the ubiquitin transfer reaction, the co-activator APC10, and one activator subunit belonging to one of two classes, respectively called CELL DIVISION CYCLE 20 (CDC20) or CDC20 HOMOLOG 1 (CDH1), the latter also known in plants as CELL CYCLE SWITCH 52 (CCS52) (Tarayre et al., 2004; Kevei et al., 2011; Heyman and De Veylder, 2012). The activator proteins recruit the APC/C ubiquitination targets through recognition of conserved amino acid motifs such as the Destruction box (D-box) (Pfleger and Kirschner, 2000; De Veylder et al., 2007; da Fonseca et al., 2011).

The plant *CCS52* gene was first identified in *Medicago*, where it plays an important role in the establishment of the polyploid tissues of the root nodules (Cebolla et al., 1999). The described link between *CCS52* expression, initiation of differentiation, and the onset of the endocycle was later confirmed in other plant species. For example, in tomato, decreased *CCS52A* levels were found to cause a reduction in endoreplication and fruit size, whereas in rice, mutation of *CCS52A* resulted in dwarf growth and problems with kernel development due to a reduction of endoreplication in the endosperm (Mathieu-Rivet et al., 2010; Su’udi et al., 2012; Xu et al., 2012).

In Arabidopsis, three isoforms of CCS52 are present, two A-types (CCS52A1 and CCS52A2) and one plant specific B-type (CCS52B) (Tarayre et al., 2004; Kevei et al., 2011). Prophase-confined expression of *CCS52B* indicates that it might play a role in the degradation of M-phase proteins necessary for the progression of mitosis (Yang et al., 2017). Contrary, the *CCS52A1* and *CCS52A2* genes are thought to repress cell division in a tissue-specific manner that is determined by their expression pattern. Within the root, *CCS52A1* is predominantly expressed at the root elongation zone where it controls cell cycle exit, illustrated by an increased root meristem size in *ccs52a1* knockout plants (Vanstraelen et al., 2009). Additionally, *CCS52A1* expression can be found in leaves and trichomes, where it controls the number of endocycles (Lammens et al., 2008; Boudolf et al., 2009; Larson-Rabin et al., 2009; Baloban et al., 2013; Heyman et al., 2017). Next to controlling endocycle progression in the leaf, CCS52A2 appears to be important for maintaining the low proliferation status of the quiescent center (QC) and the organizing center (OC) of respectively the root and the shoot, seeing that *ccs52a2-1* mutant plants show a severe disruption of meristem organization, leading to a short root, dwarf growth and a strong reduction in the development of reproductive organs (Vanstraelen et al., 2009; Liu et al., 2012).

Currently, only a relatively limited set of proteins have been thoroughly characterized as targets of the CCS52-activated APC/C. In Arabidopsis, protein stability of the A-type cyclin CYCA2;3 was found to be reduced by APC/C^CCS52A1^ to control the onset of endoreduplication (Boudolf et al., 2009). The ETHYLENE RESPONSE FACTOR 115 (ERF115) transcription factor was initially identified as an interactor of CCS52A2 in a tandem affinity purification experiment and was shown to be an important rate-limiting factor of QC cell division (Heyman et al., 2013). Another protein identified as a CCS52A2 target is Cellulose Synthase Like-D 5 (CSLD5), a cell wall biosynthesis enzyme that plays a role in the assembly of the newly forming cell plate during division, and that is rapidly degraded upon completion of mitosis, but not in the *ccs52a2-1* mutant background (Gu et al., 2016). In rice, the CCS52 homolog TILLER ENHANCER (TE) / TILLERING AND DWARF (TAD1) was shown to mediate the ubiquitination and subsequent degradation of MONOCULM 1 (MOC1), called LATERAL SUPPRESSOR (LAS) in Arabidopsis, a GRAS-family transcription factor that promotes shoot branching and tillering (Lin et al., 2012; Xu et al., 2012).

Here, we have utilized an ethyl methanesulfonate (EMS) suppressor screen to identify novel APC/C^CCS52A2^ targets, based on the growth inhibitory phenotype of *ccs52a2-1* knockout plants. We show that one of the identified revertants encodes a mutant allele of the *CYCA3;4* gene and demonstrate this cyclin to be a specific target of APC/C^CCS52A2^ to ensure correct stem cell organization.

## RESULTS

### Identification of *pkn2* as a *ccs52a2-1* Suppressor Mutant

Compared to wild type (WT, Col-0) plants, *ccs52a2-1* mutant seedlings display a short root phenotype (Figures 1A and 1B; Supplemental Figures 1A and 1B) (Vanstraelen et al., 2009; Heyman et al., 2013). This phenotype was used to screen for putative targets or regulators of the APC/C^CCS52A2^ ubiquitin ligase complex through a mutagenesis revertant screen. Therefore, ethyl methanesulfonate (EMS)-mutagenized *ccs52a2-1* plants were screened in the M_2_ generation for a recovered root growth. Out of a total of 260 initially identified revertants, 33 were confirmed in the next generation. Among these, one revertant mutation, named *pikmin 2* (*pkn2*), yielded a root length in between that of WT and *ccs52a2-1* mutant plants (Figures 1A to 1C; Supplemental Figures 1A to 1C).

**Figure 1.**
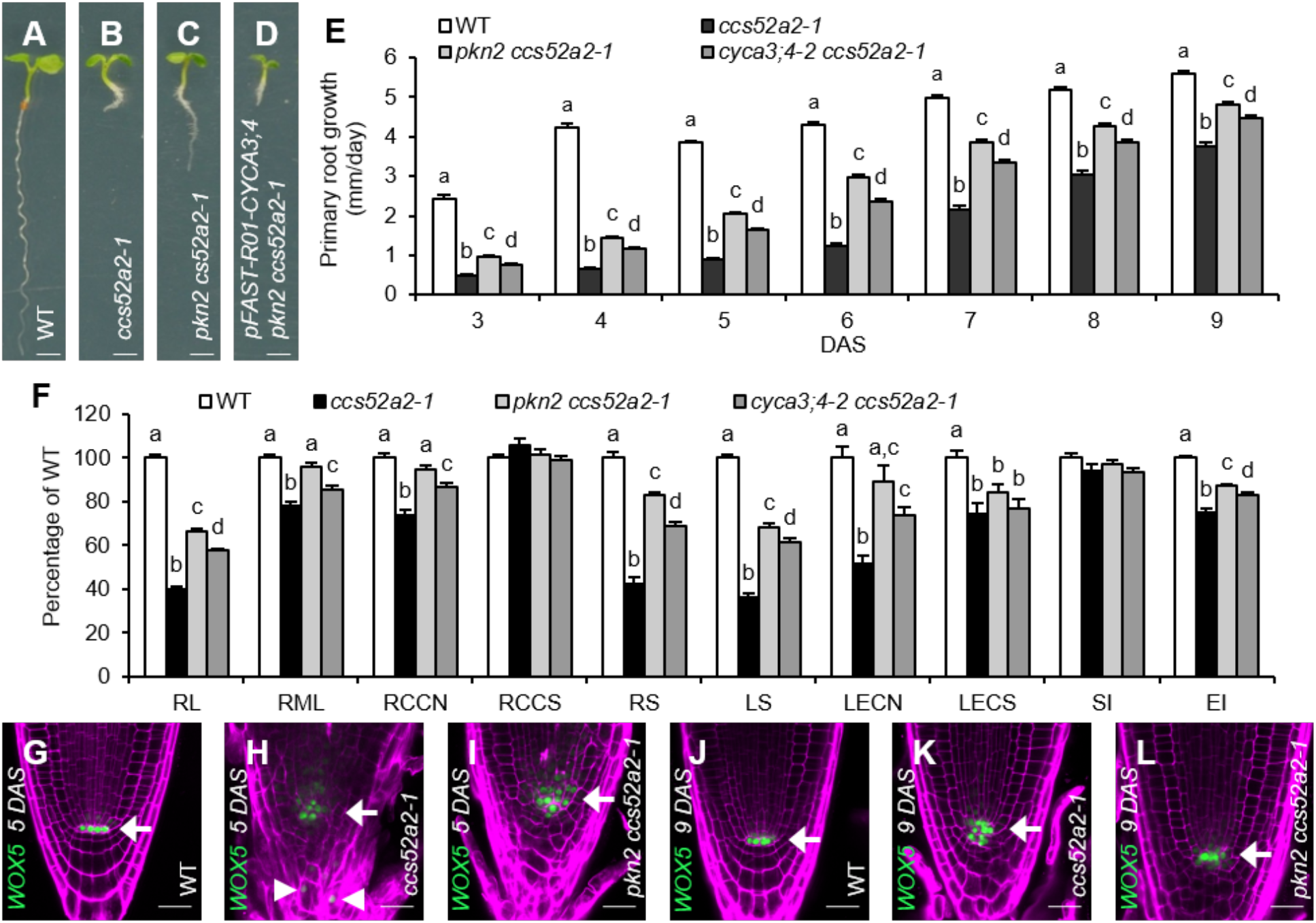
The *pkn2* Mutation Rescues the *ccs52a2-1* Short Root Phenotype and Is Located in the Cyclin Gene *CYCA3;4*. **(A-D)** Representative seedlings of WT **(A)**, *ccs52a2-1* **(B)** and *pkn2 ccs52a2-1* without **(C)** and with **(D)** the *pFAST-R01-CYCA3;4* complementation construct at 5 DAS. Scale bars represent 1 mm. **(E-F)** Growth characteristics of WT, *ccs52a2-1* and the double mutants *pkn2 ccs52a2-1* and *cyca3;4-2 ccs52a2-1*. Primary root growth from 3 to 9 DAS **(E)**. Phenotypes of the primary root at 9 DAS and the shoot and the first leaf pair at 21 DAS **(F)**. RL, root length; RML, root meristem length; RCCN, root cortical cell number; RCCS, root cortical cell size; RS, rosette size; LS, leaf size; LECN, leaf epidermal cell number; LECS, leaf epidermal cell size; SI, stomatal index; EI, endoreplication index. Error bars represent standard error (n ≥ 15). Letters on the bars indicate statistically different means (P < 0.05, mixed model analysis, Tukey correction for multiple testing). **(G-L)** Representative confocal images of *WOX5_pro_:GFP-GUS* expressing WT **(G&J)**, *ccs52a2-1* **(H&K)** and *pkn2 ccs52a2-1* **(I&L)** primary roots at 5 **(G-I)** and 9 DAS **(J-L)**. GFP signal is shown in green, while cell walls are visualized through propidium iodide staining (magenta). Arrows indicate the position of the QC, while ectopic *WOX5* expression in *ccs52a2-1* is indicated by arrowheads. Scale bars represent 25 μm.

Root growth of the *ccs52a2-1* mutant was found to be strongly reduced during early development, showing a primary root growth rate of only around 20% of that of WT plants from 3 to 5 days after stratification (DAS) (Figure 1E). At later time points, root growth of the *ccs52a2-1* mutant gradually recovered, but never fully caught up to that of WT plants. At 9 DAS, the *ccs52a2-1* root length was about 40% of that of WT plants (Figure 1F). Compared to the *ccs52a2-1* mutant, the *pkn2 ccs52a2-1* double mutant showed an increased root growth rate over the total time frame measured (Figure 1E), resulting in a root length recovery to 67% of that of WT plants at 9 DAS (Figure 1F). The root growth phenotype of *ccs52a2-1* was reflected in the root meristem length measured at 9 DAS, reaching only 78% of wild type, primarily caused by a reduction in cell number as cell size was not significantly altered (Figure 1F). In the *pkn2 ccs52a2-1* double mutant, root meristem length and cell number were slightly smaller but not significantly different from WT plants, nor was the cortical cell size (Figure 1F).

A striking characteristic of the *ccs52a2-1* mutant is a disorganized root stem cell niche, due to a loss of QC cell quiescence (Vanstraelen et al., 2009). To examine this phenotype in detail, a *WOX5_pro_:GFP-GUS* transcriptional reporter that marks the QC cells was introgressed into the *ccs52a2-1* and *pkn2 ccs52a2-1* mutant backgrounds. During early development (at 5 DAS), *WOX5* expression was detected in an expanded area of the disorganized QC and stem cell niche of the *ccs52a2-1* mutant, as well as in differentiated tissues such as the columella cells (Figures 1G and 1H; Supplemental Figures 1D and 1E). At a later developmental stage (9 DAS), *WOX5* expression was confined to the stem cell niche, coinciding with the partially recovered root growth phenotype, but still revealed a disorganized cell patterning (Figures 1J and 1K). Compared to the *ccs52a2-1* mutant, the *pkn2 ccs52a2-1* double mutant showed a slightly improved meristem organization at 5 DAS, together with a more confined *WOX5* expression domain (Figures 1H and 1I; Supplemental Figures 1E and 1F). At 9 DAS, its *WOX5* expression pattern more closely resembled that of WT plants (Figures 1J to 1L).

For the shoot tissue, a recovery of the *ccs52a2-1* phenotypes was seen in the *pkn2 ccs52a2-1* double mutant for the majority of parameters analyzed (Figure 1F). Projected rosette size of *ccs52a2-1* at 21 DAS was only 43% of that of WT plants, whereas that of the double mutant reached 83% (Figure 1F). This was reflected by the size of the first leaf pair at 21 DAS, with *ccs52a2-1* and *pkn2 ccs52a2-1* reaching 36% and 68% of WT leaf size, respectively (Figure 1F). Leaf growth recovery appeared to be mostly driven at the cell number level, with *ccs52a2-1* showing a reduction to 52% of WT epidermal cell number, whereas the *pkn2 ccs52a2-1* double mutant reached 89% (Figure 1F). No statistically significant recovery was seen in the epidermal cell size, with *ccs52a2-1* and *pkn2 ccs52a2-1* showing a similar reduction to 75% and 84% of that of WT, respectively (Figure 1F). Furthermore, neither *ccs52a2-1* nor the double mutant showed a significant change in pavement versus stomatal cell ratio, as represented by the stomatal index (Figure 1F). As previously reported (Baloban et al., 2013), the number of endocycles, as represented by the endoreplication index, was reduced in the *ccs52a2-1* mutant to 75% of that of WT plants. A moderate recovery could be observed for the *pkn2 ccs52a2-1* double mutant, with an endoreplication index of 87% of that of WT plants (Figure 1F).

### Identification of *cyca3;4* as *pkn2*

To identify the affected gene underlying the *pkn2* mutation, a mapping scheme was set up, in which the *pkn2 ccs52a2-1* mutant was backcrossed to the original *ccs52a2-1* parental line and subsequently self-pollinated. In the resulting segregating F2 mapping population, plants with the revertant long root phenotype were selected and pooled for gene mapping through next-generation sequencing, using the EMS-generated single-nucleotide polymorphisms (SNPs) as *de novo* mapping markers (See Methods for details). Plotting the distribution of the SNPs on the genome revealed a broad peak of increased mutant allele frequency in the middle of chromosome 1 and subsequently an interval of 4 million base pairs (Mbp) was selected for detailed analysis (from 13 Mbp to 18 Mbp; Supplemental Figure 2 and Supplemental Table 1). After filtering for EMS-specific mutations with a concordance above 0.8 and filtering out intergenic or intronic mutations, only one candidate gene remained, namely *AT1G47230*, encoding the A-type cyclin CYCA3;4. The identified mutation was found to be located on the acceptor splice site of intron 5, causing the acceptor G base to become an A (Figure 2A). Correspondingly, isolation of *CYCA3;4* transcript amplicons through RT-PCR identified two novel and distinct *CYCA3;4* splice variants within *pkn2 ccs52a2-1* (Figure 2B). Sequencing of these transcripts revealed that the longer variant retained the intron preceding the splice acceptor site mutation, while the shorter variant had both intron 5 plus 13 bp from the following exon spliced out (Figure 2A, Supplemental Figure 3). Cyclins generally contain two conserved cyclin folds, with the N-terminal fold responsible for binding to a CDK protein, and the C-terminal fold responsible for target binding. Both expressed mRNA variants of *pkn2* resulted in a frame shift, leading to a premature stop-codon and the loss of the second half of the predicted C-terminal cyclin fold of CYCA3;4 (Figure 2A), strongly suggesting that the mutant CYCA3;4 variants can no longer bind target proteins or perform their function.

**Figure 2.**
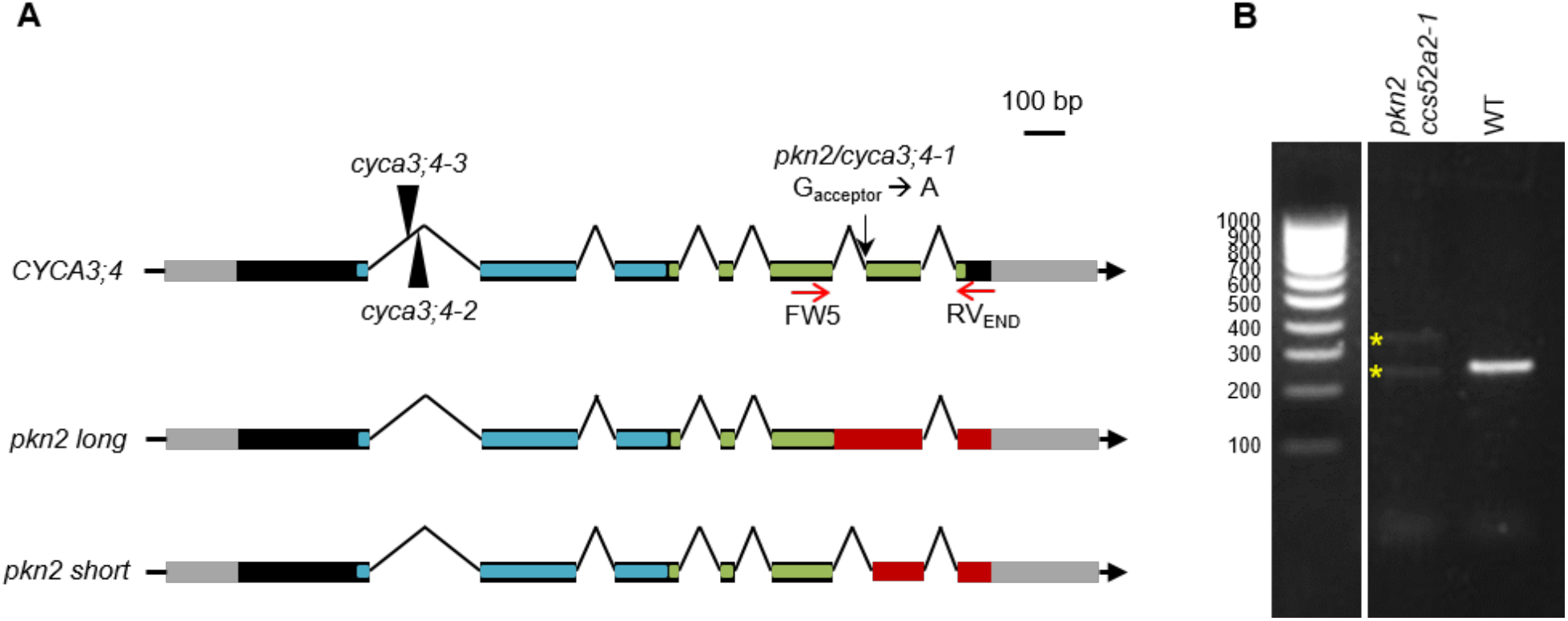
The *CYCA3;4* Gene Structure. **(A)** The gene structure of the WT *CYCA3;4* gene, showing the location of the EMS mutation (black arrow) and T-DNA insertions (black triangles), and the two splice variants created through the *pkn2* mutation. The gray and black boxes represent the untranslated regions and the coding sequences respectively, while the lines represent the intergenic sequences and introns. The predicted N- and C-terminal cyclin folds are indicated in blue and green, respectively. In the mutant variants, the out-of-frame sequence is indicated in red. **(B)** RT-PCR on whole seedling cDNA of *pkn2 ccs52a2-1* and WT (Col-0) using *CYCA3;4* primers FW5 and RVEND (represented by red arrows in **A**), and, whereas only one amplicon was detected for WT, two distinct amplicons were detected for the revertant (yellow stars), representing two newly created splice variants.

Transformation of a complementation construct containing a functional copy of the *CYCA3;4* gene into the *pkn2 ccs52a2-1* mutant confirmed that the *pkn2* mutation in *CYCA3;4* was responsible for the recovery of the *ccs52a2-1* root growth phenotype, as out of the 135 transformants obtained, 123 reverted to the stunted root growth phenotype (Figure 1D). Remarkably, many transformants grew even worse than *ccs52a2-1* plants, suggesting that the root growth phenotype of the *ccs52a2-1* plants might be strongly correlated with CYCA3;4 abundance and that timely breakdown of CYCA3;4 might be essential for proper plant development.

As independent proof that *CYCA3;4* deficiency rescues the *ccs52a2-1* phenotype, independent *CYCA3;4* mutants obtained from insertion collections were selected. Two lines, SALK_204206 and SALK_061456, named *cyca3;4-2* and *cyca3;4-3* in accordance with regarding *pkn2* as *cyca3;4-1*, were found to hold a T-DNA insertion within the first intron of the *CYCA3;4* gene that resulted in a very strong reduction of transcript abundance (Figure 2A; Supplemental Figures 4A and 4B). Both T-DNA insertion mutants were analyzed over different root and shoot parameters, but did not show any significant phenotypic differences from WT plants (Supplemental Figure 4C). However, when the *cyca3;4-2* mutant was introgressed into the *ccs52a2-1* background, the resulting *cyca3;4-2 ccs52a2-1* double mutant largely phenocopied the *pkn2 ccs52a2-1* double mutant in respect to the measured root and leaf growth parameters (Figures 1 E and 1F), displaying a partial recovery of the *ccs52a2-1* root length and meristem size, leaf size, leaf epidermal cell number, and endoreduplication phenotypes, albeit slightly less than compared to *pkn2 ccs52a2-1*.

As *CYCA3;4* is part of a four-member family of closely related *CYCA3* genes, it was tested whether the absence of another family member shared the capacity to rescue the *ccs52a2-1* mutant phenotype. For this, an available homozygous *CYCA3;1* T-DNA insertion line (Takahashi et al., 2010) was crossed with the homozygous *ccs52a2-1* mutant. Plants from the segregating F2 population were genotyped and their root length measured. Similar to the two *CYCA3;4* insertion mutants, no root growth phenotype was observed for the *CYCA3;1* mutant (Supplemental Figure 5). Moreover, contrary to what was observed for *CYCA3;4*, a lack of *CYCA3;1* did not result in a rescue of the *ccs52a2-1* short root phenotype (Supplemental Figure 5).

### CCS52A2 Targets CYCA3;4 for Degradation

CYCA3;4 is likely to be a direct target for APC/C^CCS52A2^-dependent ubiquitination and subsequent proteasomal degradation, as it holds the full D-box sequence RVVLGELPN, which serves as a binding site recognition motif for the APC/C (da Fonseca et al., 2011). To test this hypothesis, a previously described *CYCA3;4_pro_:CYCA3;4-GUS* translational reporter line(Bulankova et al., 2013) was treated with the proteasome inhibitor MG-132. For comparison, the corresponding translational reporter of *CYCA3;1* (*CYCA3;1_pro_:CYCA3;1-GUS*) was included. Following a short GUS staining, CYCA3;1-GUS activity was barely visible in the root tip, whereas CYCA3;4-GUS could be detected in the root elongation zone (Figures 3A and 3C). After a 24-h treatment with MG-132, an increase in GUS activity could be observed in the root elongation zone for CYCA3;1-GUS (Figure 3B). This increase was more outspoken for CYCA3;4-GUS, showing stronger staining in not only the elongation zone, but also the root meristem (Figure 3D). The accumulation of CYCA3;4-GUS in the root meristem corresponded to the spatial expression pattern of *CCS52A2*, whereas expression of *CCS52A1* was confined to the root elongation zone (Supplemental Figure 6) (Vanstraelen et al., 2009; Liu et al., 2012). These data suggest that CYCA3;1 might rather be targeted for degradation by CCS52A1, whereas CYCA3;4 might be controlled by both APC/C^CCS52A1^ and APC/C^CCS52A2^.

**Figure 3.**
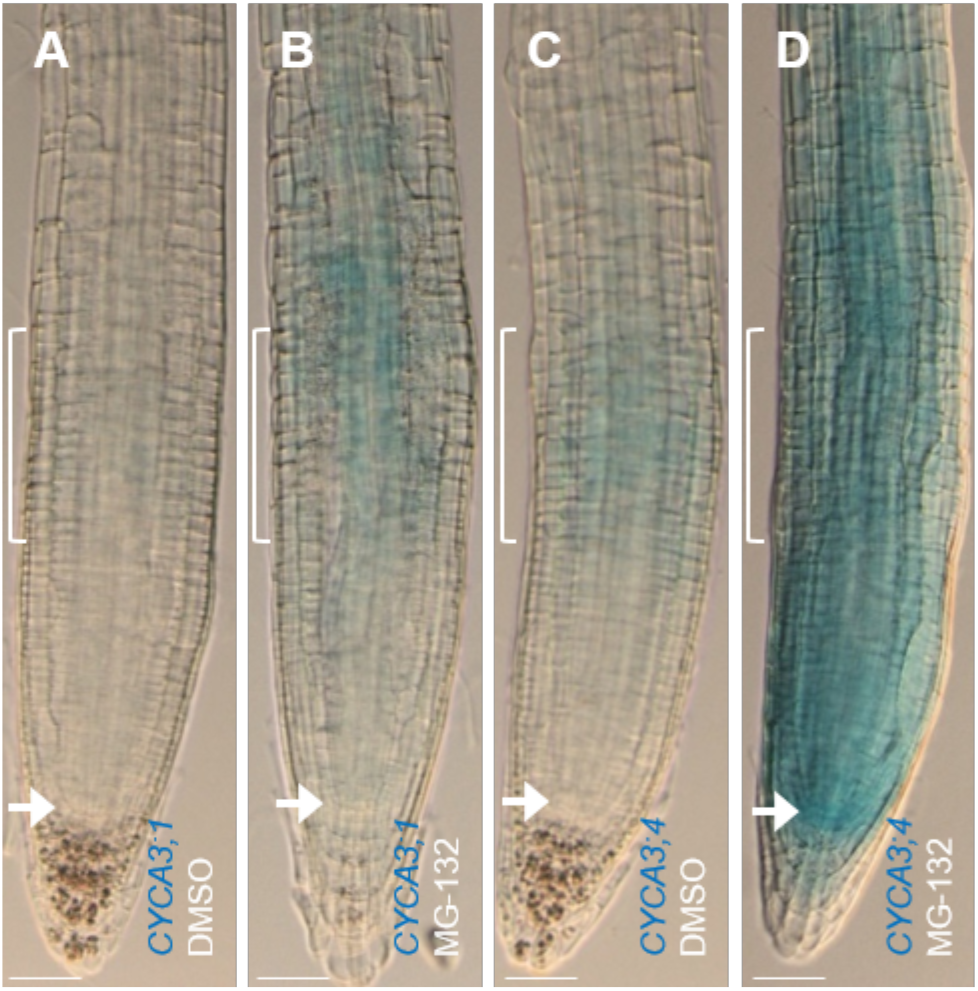
CYCA3 Protein Levels Are Dependent on Proteasomal Degradation. Histochemical GUS staining of *CYCA3;1_pro_:CYCA3;1-GUS* **(A-B)** and *CYCA3;4_pro_:CYCA3;4-GUS* **(C-D)** root tips at 5 DAS after 24-h treatment with DMSO control **(A and C)** or with the proteasome blocker MG-132 **(B and D)**. The transition zone and the QC are indicated by brackets and arrows, respectively. Roots were stained for 30 min. Scale bars represent 50 μm.

To confirm the hypothesis that CYCA3;4 is marked for breakdown by CCS52A2 in the root meristem, the *CYCA3;4_pro_:CYCA3;4-GUS* and *CYCA3;1_pro_:CYCA3;1-GUS* reporters were introgressed into the *ccs52a2-1* mutant background. Strikingly, plants mutant for *CCS52A2* and homozygous for the *CYCA3;4_pro_:CYCA3;4-GUS* construct showed severe root growth reduction compared to *ccs52a2-1* plants without the construct, while *CYCA3;1_pro_:CYCA3;1-GUS* did not exert such effect (Figures 4A to 4D, and 4I). This is most probably due to the extra CYCA3;4 protein present because of the reporter construct, again highlighting the importance of a timely breakdown of CYCA3;4 for plant development. Comparing the GUS activity of the *CYCA3;1_pro_:CYCA3;1-GUS* and *CYCA3;4_pro_:CYCA3;4-GUS* constructs in WT versus *ccs52a2-1* mutant plants revealed that the spatial accumulation pattern of CYCA3;1-GUS was largely maintained in the shortened *ccs52a2-1* meristem (Figures 4 E and 4F). Contrastingly, CYCA3;4-GUS staining appeared to concentrate at the most distal part of the root meristem (Figures 4G and 4H).

**Figure 4.**
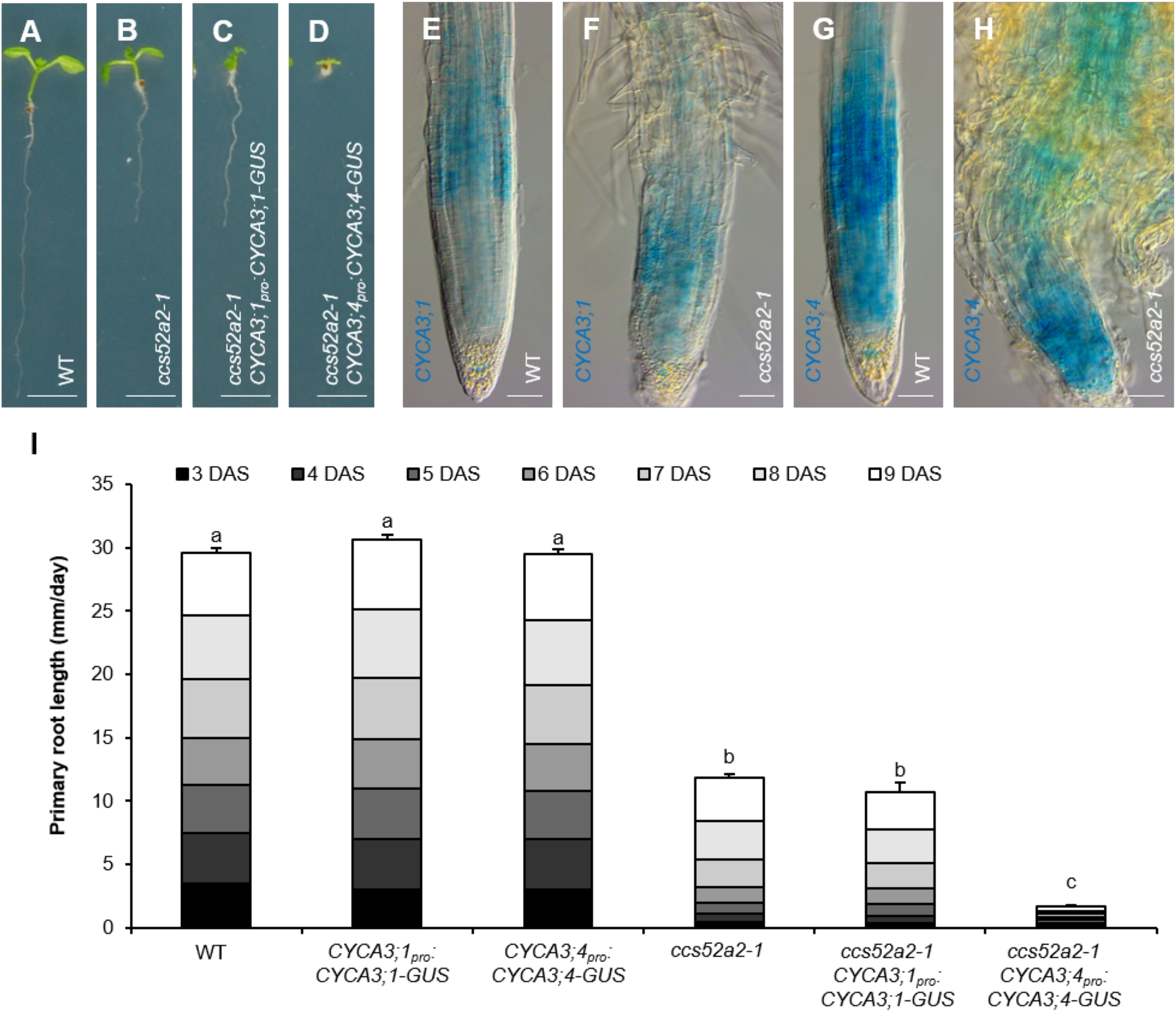
The *CYCA3_pro_:CYCA3-GUS* Constructs in the *ccs52a2-1* Background. **(A-D)** Representative seedlings of WT **(A)**, *ccs52a2-1* **(B)** and *ccs52a2-1* with *CYCA3;1_pro_:CYCA3;1-GUS* **(C)** or *CYCA3;4_pro_:CYCA3;4-GUS* **(D)** at 9 DAS. Scale bars represent 5 mm. **(E-H)** Histochemical GUS staining at 5 DAS of WT **(E and G)** and *ccs52a2-1* KO **(F and H)** root tips with either *CYCA3;1_pro_:CYCA3;1-GUS* **(E-F)** or *CYCA3;4_pro_:CYCA3;4-GUS* **(G-H)** constructs in their background. Roots were stained for 2 h. Pictures were taken at the same magnification. Scale bars represent 50 μm. **(I)** Primary root length of the respective *CYCA3_pro_:CYCA3-GUS* lines in the *ccs52a2-1* background from 3 to 9 DAS. Error bars represent standard error (n ≥ 25), and letters indicate statistically different growth for each genotype (P < 0.0001, mixed model analysis, Tukey correction for multiple testing).

To identify the cell cycle phase during which both cyclins might be targeted for destruction, root tips of the *CYCA3* translational reporter lines in WT background were arrested at different points in the cell cycle using hydroxyurea (HU), bleomycin or oryzalin, arresting the cell cycle in the S-, G_2_- or early M-phase, respectively. Compared to control conditions, an increased GUS staining could be seen after HU treatment for CYCA3;1-GUS, most prominently in the elongation zone, whereas a decrease was observed for CYCA3;4-GUS throughout the root tip (Figure 5). Conversely, the CYCA3;1-GUS staining pattern and intensity following bleomycin or oryzalin did not differ from the control, whereas the CYCA3;4-GUS signal in the root meristem was strongly increased under those conditions (Figure 5). These data indicate that the CYCA3;1 and CYCA3;4 proteins are differently controlled in both a temporal and spatial manner, with CYCA3;1 being predominantly present during the S-phase, while CYCA3;4 appears to accumulate during the G_2_- or early M-phase.

**Figure 5.**
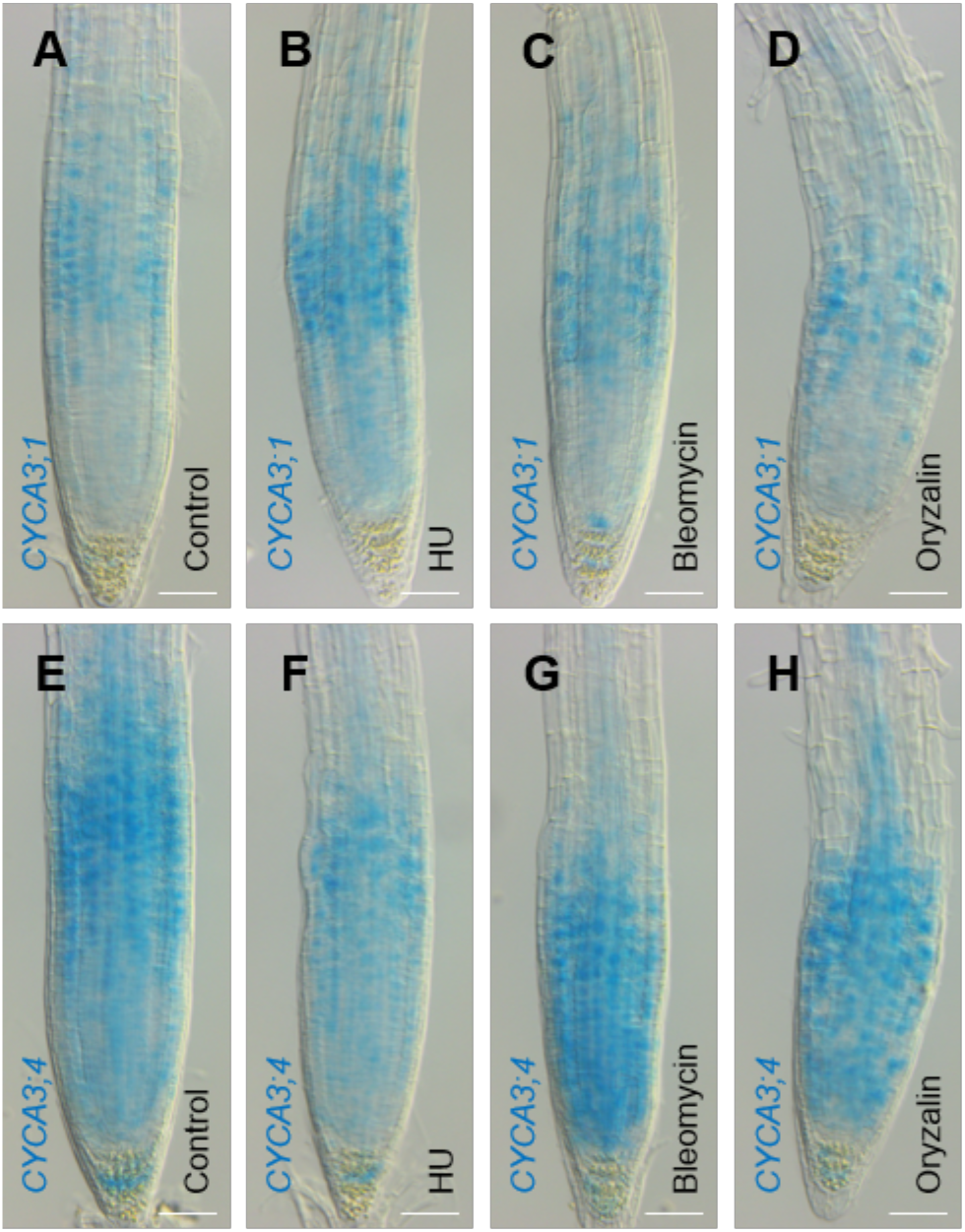
Expression of CYCA3 Proteins Throughout the Cell Cycle. Root tip GUS signal of *CYCA3;1_pro_:CYCA3;1-GUS* **(A)** or *CYCA3;4_pro_:CYCA3;4-GUS* **(B)** at 6 DAS, under control conditions or treated with hydroxyurea (HU), bleomycin or oryzalin, respectively blocking the cell cycle in S-, G_2_- or M-phase. Scale bars represent 50 μm.

To pinpoint more precisely the cell cycle phase during which CYCA3;4 would be targeted for destruction by APC/C^CCS52A2^, the CYCA3;4-GUS protein abundance in the root tip was analyzed through an immunostaining experiment using an anti-GUS antibody. In WT plants, a positive CYCA3;4-GUS signal could only be detected in nuclei of prophase cells (Figure 6A, Supplemental Table 2). By contrast, in the *ccs52a2-1* mutant background, CYCA3;4-GUS could additionally be detected in metaphase and anaphase nuclei (Figure 6B, Supplemental Table 2), demonstrating that CYCA3;4 is targeted for destruction by APC/C^CCS52A2^ in post-prophase cells.

**Figure 6.**
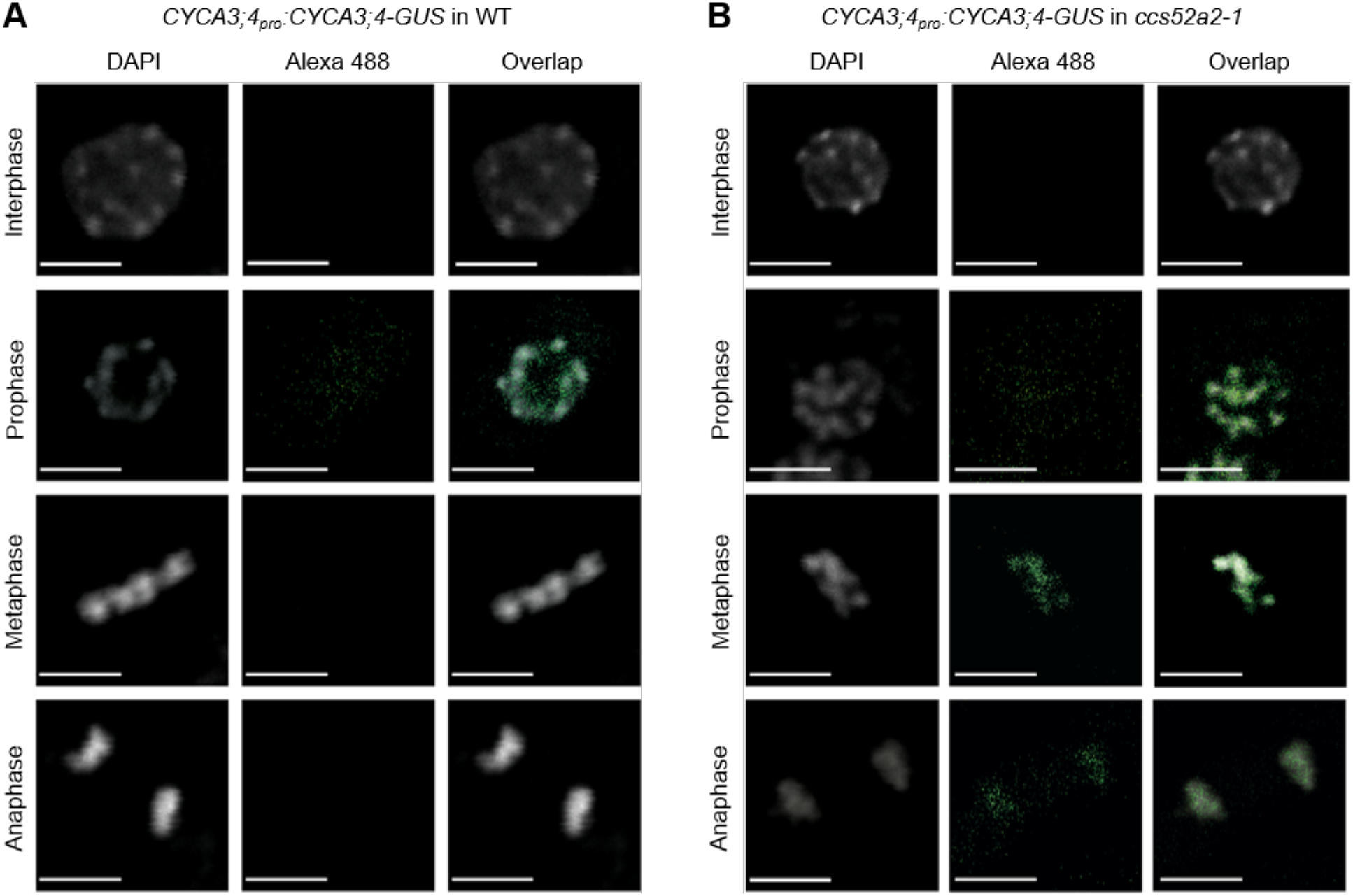
The Accumulation of CYCA3;4 Persists Past Prophase in the *ccs52a2-1* Background. Immunostaining of CYCA3;4-GUS throughout the cell cycle in squashed root tips of plants containing the *CYCA3;4_pro_:CYCA3;4-GUS* construct in the WT **(A)** or *ccs52a2-1* **(B)** background. DNA was stained using DAPI (gray) and CYCA3;4-GUS was visualized with a polyclonal rabbit anti-GUS primary antibody and an Alexa-488 secondary antibody (green). Scale bars represent 5 μm.

### Moderate *CYCA3;4* Overexpression Induces Unscheduled Formative Divisions in the Root Meristem, Whereas High Overexpression Inhibits Cell Division

The data suggested that CYCA3;4 abundance needs to be strictly controlled, as its stabilization appears to trigger a growth arrest. Therefore, to study the effects of increased CYCA3;4 levels in more detail, overexpression lines were generated expressing the *CYCA3;4* gene from the strong *Cauliflower Mosaic Virus 35S* promoter (*CYCA3;4^OE^*). Overexpression levels in the root tip ranged between two- to eightfold compared to WT levels, whereas in the young shoot relative overexpression levels were higher, varying between 16- to 29-fold (Supplemental Figure 7A). Homozygous plants were generally smaller but appeared to be prone to tissue- and development-dependent silencing of the overexpression construct, as evidenced by the difference in penetrance of the observed phenotypes (Supplemental Figure 7B). This silencing could be reverted by crossing with WT plants and generating hemizygous lines. Therefore, to be able to see the effect of both moderate and high levels of overexpression, analysis in the root was performed on homozygous lines 11.2 and 12.4, which showed partial silencing of the overexpression construct, as well as on hemizygous plants resulting from crossing the respective lines with WT plants. Initially, following germination, root growth in both the homozygous and hemizygous *CYCA3;4^OE^* lines was similar to WT plants, but subsequently became slower, most prominently observed in the hemizygous lines, resulting in a significant reduction in total root length at 9 DAS (Figure 7A). This reduced growth correlated with a decrease in root meristem length, being about 20% shorter for both homozygous lines and around 40% for both hemizygous lines (Figure 7B). This shortening was found to be mostly due to a decrease in meristem cell number, while cell size remained relatively unchanged (Figure 7B). Interestingly, an aberrant division pattern reminiscent to that of *ccs52a2-1* mutant roots could be detected in the majority of the measured roots of the homozygous lines, whereas the cell pattern in the highly overexpressing hemizygous lines appeared normal (Figure 7C to 7E). Taken together, these data indicate that moderate overexpression of *CYCA3;4* induces unscheduled formative divisions, whereas high overexpression inhibits cell division all together.

**Figure 7.**
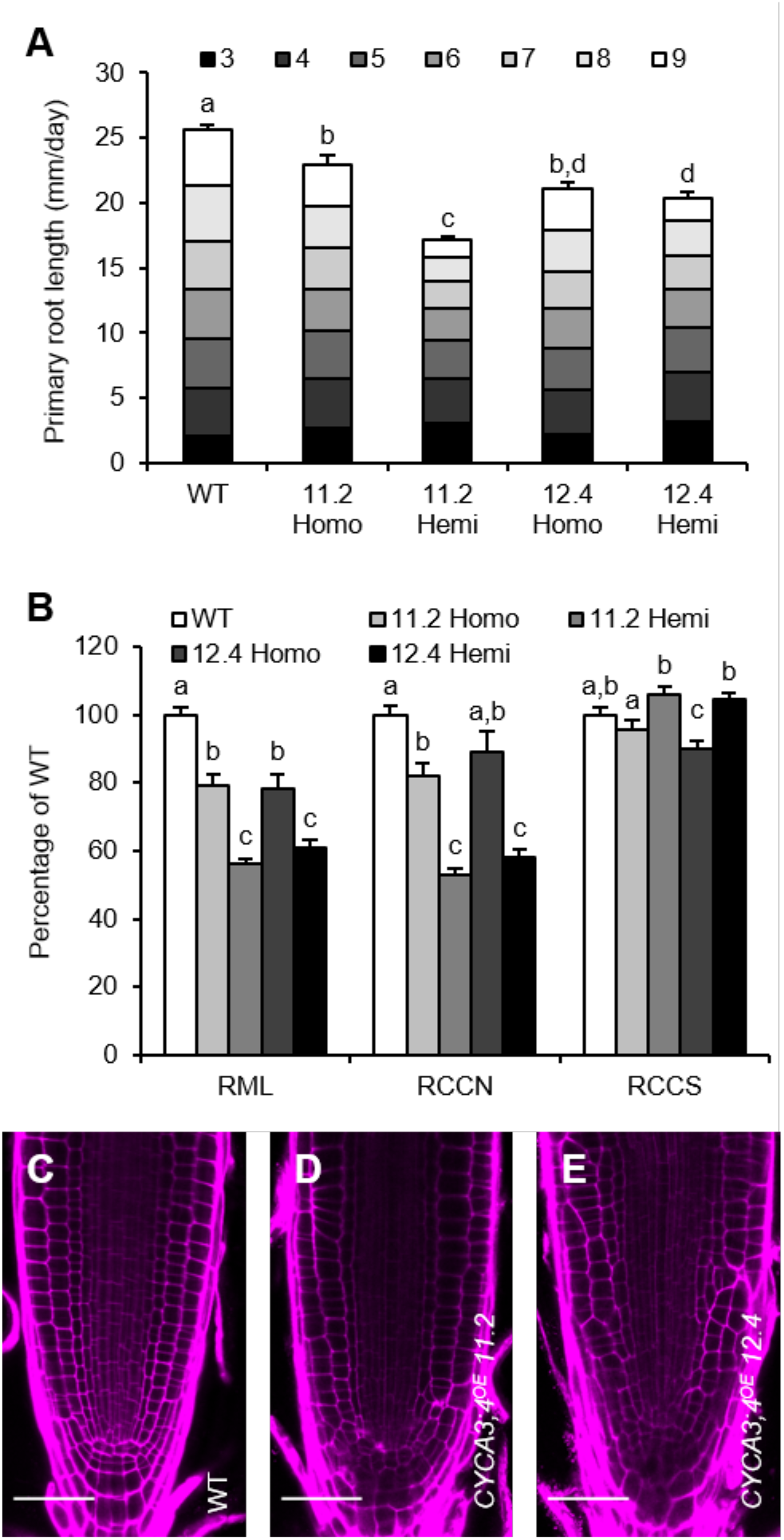
The Effects of *CYCA3;4* Overexpression in the Root. **(A-B)** Phenotypical analysis of homozygous and hemizygous *CYCA3;4^OE^* lines 11.2 and 12.4. Primary root length from 3 to 9 DAS **(A)**; and RML, root meristem length; RCCN, root cortical cell number; and RCCS, root cortical cell size at 9 DAS **(B)**. Error bars represent standard error (n ≥ 9). Letters represent significantly different means (P < 0.05, mixed model analysis, Tukey correction for multiple testing). **(C-E)** Representative confocal images of the root meristem of WT **(C)** and homozygous *CYCA3;4^OE^* lines 11.2 **(D)** and 12.4 **(E)**. Cell walls were stained using propidium iodide staining. Scale bars represent 25 μm.

### *CYCA3;4* Overexpression Severely Impacts Leaf Cell Differentiation

Although homozygous *CYCA3;4^OE^* lines 11.2 and 12.4 showed a strongly reduced root meristem size, the size of the first leaf pair was only slightly reduced, whereas that of other independent lines was strongly affected, indicating age-dependent silencing of the overexpression construct (Supplemental Figure 7B). Therefore, we focused on the strongly overexpressing *CYCA3;4^OE^* hemizygous lines 11.2 and 12.4 for leaf phenotyping, in which maintenance of *CYCA3;4* overexpression was confirmed through qRT-PCR (Supplemental Figure 7C). The size of both the whole rosette as well as the first leaf pair was dramatically reduced within both independent lines to less than 20% of that of WT plants (Figure 8A). This reduction was due to a lack of cell growth, as pavement cells were round and small (Figures 8B to 8D), with their size reduced to only about 5% of that of WT cells (Figure 8A). Concurrently, endoreplication was also strongly suppressed in both hemizygous lines (Figure 8A). Interestingly, while pavement cell number was increased (Figure 8A), an almost complete lack of stomata could be observed. Likewise, a less severe but significant reduction in stomatal density was observed in all homozygous lines (Supplemental Figure 7B). The observed reduction in stomatal number was accompanied by an increase in transcripts of genes controlling the early steps of stomata formation, including *SPCH, SDD1* and *MUTE* (Figure 8H). Contrary, the expression of the late stomata pathway gene *FAMA* was not significantly altered (Figure 8H).

**Figure 8.**
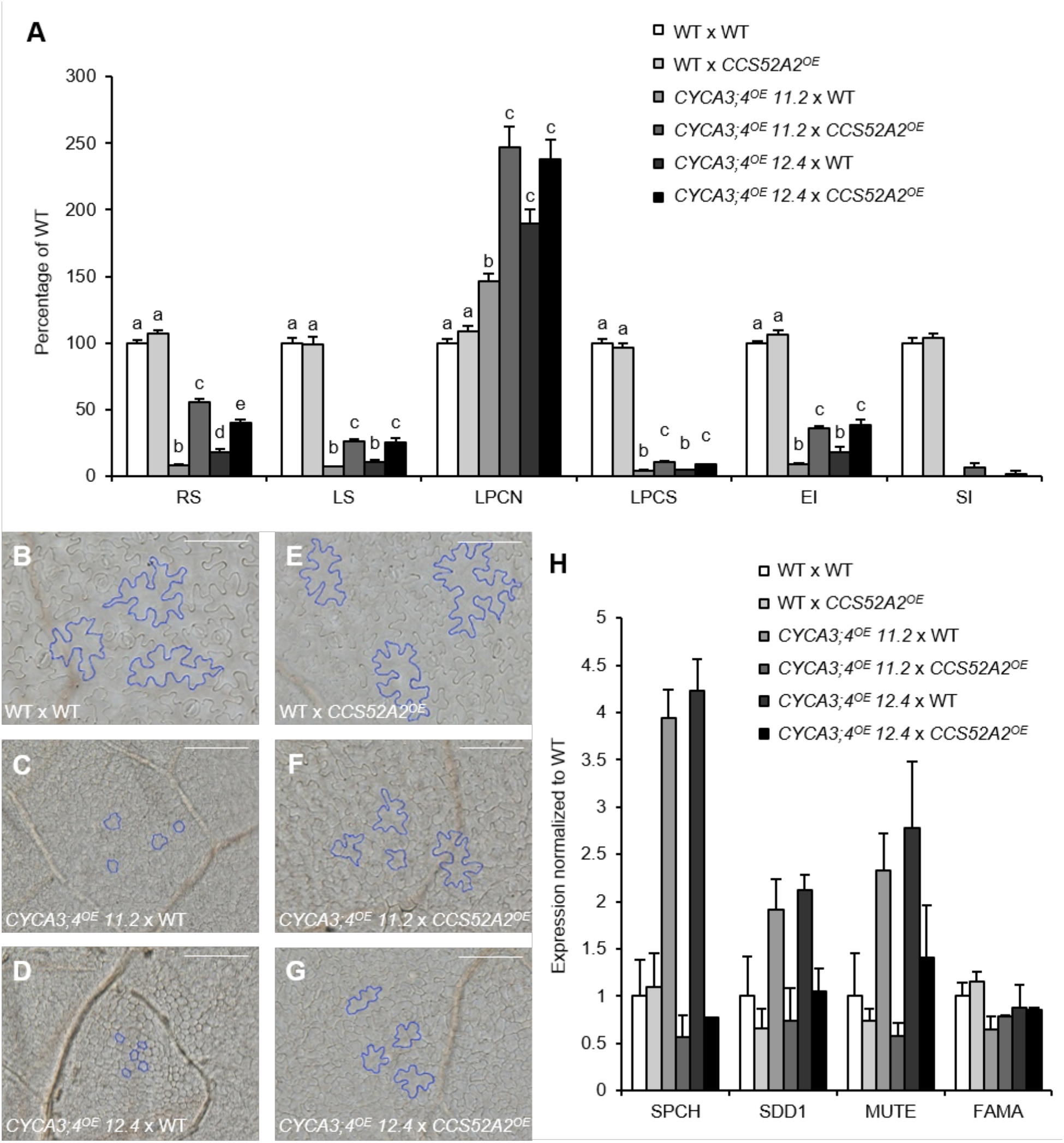
The Effects of Overexpression of *CYCA3;4* and Co-Overexpression of *CCS52A2* in the Leaf. **(A)** Shoot phenotyping at 21 DAS of hemizygous first-generation progeny resulting from crosses between WT, *CYCA3;4^OE^* line 11.2 or line 12.4 and WT or *CCS52A2^OE^*. RS, rosette size; LS, leaf size of the first leaf pair; LPCN, leaf pavement cell number; LPCS, leaf pavement cell size; EI, endoreplication index. Error bars represent standard error (n ≥ 5). Letters on the bars indicate statistically different means (P < 0.05, mixed model analysis, Tukey correction for multiple testing). **(B-G)** Pictures of the leaf epidermis, with some cells highlighted in blue to emphasize the change in cell size and shape, of the following crosses: WT x WT **(B)**, *CYCA3;4^OE^ 11.2* x WT **(C)**, *CYCA3;4^OE^ 12.4* x WT **(D)**, WT x *CCS52A2^OE^* **(E)**, *CYCA3;4^OE^ 11.2* x *CCS52A2^OE^* **(F)**, *CYCA3;4^OE^ 12.4* x *CCS52A2^OE^* **(G)**. Scale bars represent 50 μm. **(H)** Expression levels of the stomatal development pathway genes SPCH, SDD1, MUTE and FAMA as measured by qRT-PCR in the first leaf pair at 21 DAS. Error bars represent standard error (n = 2 or 3).

### Ectopic *CCS52A2* Expression Partially Counteracts the Leaf *CYCA3;4* Overexpression Phenotypes

Following the hypothesis that CYCA3;4 is targeted for proteasomal degradation by APC/C^CCS52A2^, it could be reasoned that the *CYCA3;4^OE^* phenotypes could be counteracted by co-overexpression of *CCS52A2*. To test this, *CYCA3;4^OE^* lines 11.2 and 12.4 were crossed with a mild *CCS52A2^OE^* line (Baloban et al., 2013) and growth characteristics were subsequently analyzed in the first-generation progeny. To rule out the effect of silencing on *CYCA3;4* transcript overabundance, overexpression of *CYCA3;4* and *CCS52A2* was confirmed by qRT-PCR (Supplemental Figure 7B). The *CYCA3;4^OE^ CCS52A2^OE^* co-overexpressing lines showed a significant recovery of growth compared to single *CYCA3;4^OE^*, as seen in rosette growth, first leaf pair size, and endoreplication index (Figure 8A). The increase in leaf size was due to an increase in pavement cell number compared to single *CYCA3;4^OE^* plants and a simultaneous increase in pavement cell size, showing again a more puzzle piece-like shape (Figures 8A and 8E to 8G). Additionally, although still limited in number, stomatal guard cells could be observed, accompanied by a normalization of transcript levels of stomatal lineage genes (Figure 8H). These results indicate that the growth recovery seen in double overexpressing plants is due to an increased targeting of the overabundant CYCA3;4 protein for proteasomal degradation by the APC/C^CCS52A2^.

### CYCA3;4 Might Function Through RBR1 Phosphorylation

In order to identify potential targets for CYCA3;4-dependent CDK phosphorylation, a phosphoproteomics assay to discover differentially phosphorylated proteins was performed on three pools of 14-day-old seedlings of the hemizygous *CYCA3;4^OE^* line 11.2. A total of 56 differentially phosphorylated peptides were identified among 54 different proteins, of which 17 phosphopeptides from 16 proteins showed enhanced phosphorylation in the *CYCA3;4^OE^* background compared to WT, whereas 39 phosphopeptides from 38 proteins displayed reduced phosphorylation (Figure 9A; Supplemental Data Sets 1 and 2). Furthermore, 28 phosphopeptides from 24 proteins were identified in only one genotype and were designated “unique”, with 19 phosphopeptides from 15 proteins only identified in WT plants and nine only identified in the *CYCA3;4^OE^* background (Supplemental Data Set 3). Interestingly, 22 of the 26 phosphopeptides (i.e. 84.6 %) being more abundantly phosphorylated in or unique for *CYCA3;4^OE^*, contained the minimal CDK phosphorylation sites SP or TP (in short [S/T]P) and out of those, four were part of the full CDK phosphorylation site [S/T]Px[K/R] (Figure 9B).

**Figure 9.**
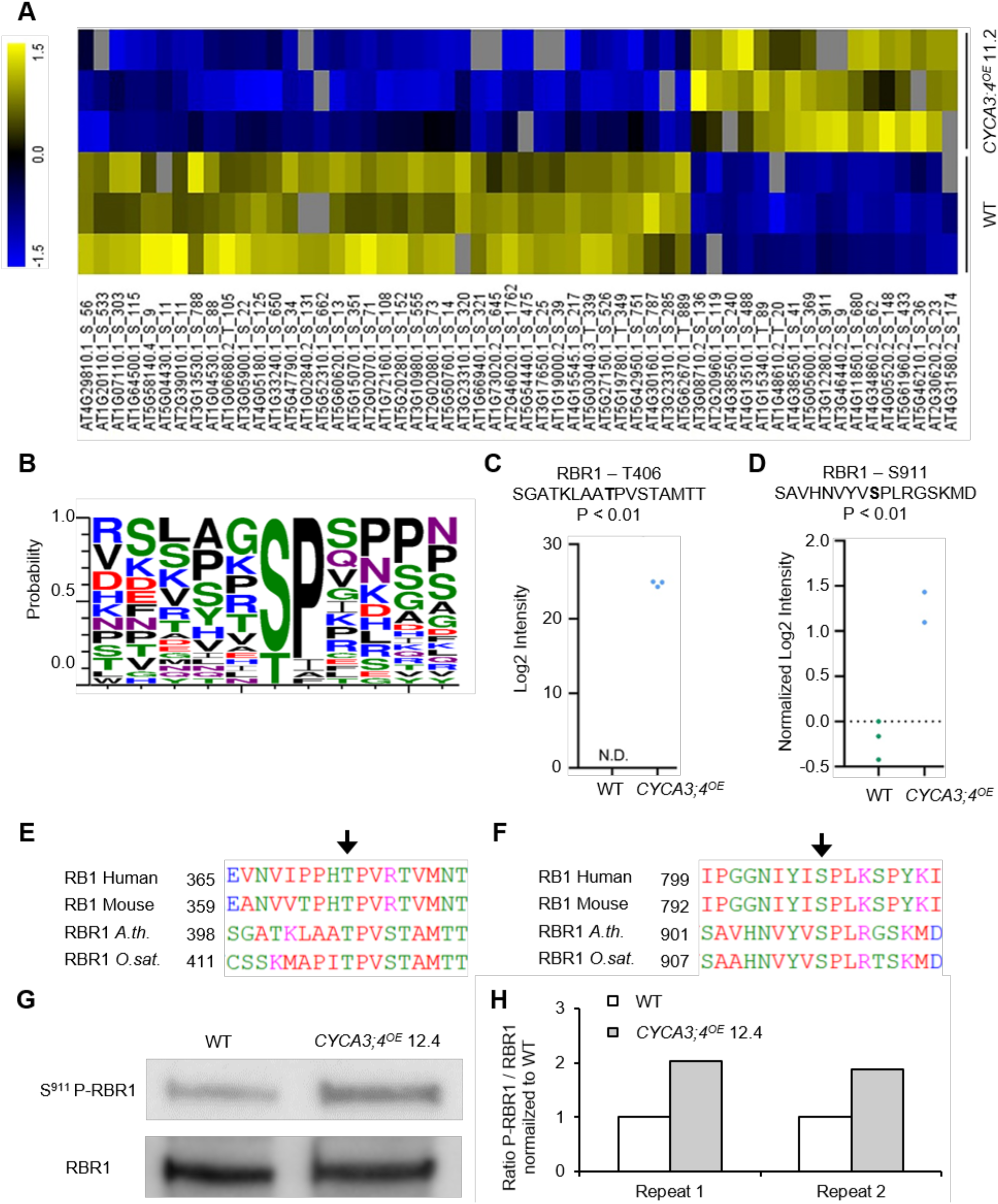
Phosphoproteomics Analysis in the *CYCA3;4^OE^* Background. **(A)** Clustering of the differentially phosphorylated sites identified. **(B)** Motif representing the occurrence of different amino acids in a ±5 amino acid interval around the phosphorylated serine or threonine in those sites showing increased phosphorylation in the *CYCA3;4^OE^* background. Picture was made using the website http://weblogo.threeplusone.com/. **(C-D)** Levels for the indicated RBR1 phosphopeptides in the WT and *CYCA3;4^OE^* phosphoproteomes. Each dot represents a biological replicate. N.D., not detected. **(E-F)** Conservation in plants and animals of phosphorylated sites Thr406 (**E**) and Ser911 (**F**) identified in *Arabiopsis thaliana* RBR1. Homologous proteins were identified using the STRING database (www.string-db.org) and alignment was performed using the CLUSTAL OMEGA web tool (https://www.ebi.ac.uk/Tools/msa/clustalo/). **(G)** Western blot of Ser911 phospho-RBR1 and RBR1, showing an increased amount of phosphorylated RBR1 in the *CYCA3;4*-overexpressing background. **(H)** Quantification of the western blot shown in **G** and one additional repeat, ratio of S911-phosphorylated RBR1 over unphosphorylated RBR1, normalized to WT.

Among the proteins showing increased phosphorylation, Histone 1.2 (H1.2, AT2G30620) and RETINOBLASTOMA-RELATED 1 (RBR1) could be found. For the latter, two CDK phosphorylation consensus sites were identified: Thr406 phosphorylation was uniquely found in the overexpression background, whereas Ser911 was 2.75 times more phosphorylated in the *CYCA3;4^OE^* background compared to the WT (Figures 9C and 9D). Both sites are highly conserved throughout the plant and animal kingdoms, with Thr406 and Ser911, respectively, being part of a conserved TP and SPx[K/R] site (Figures 9E and 9F). To confirm the increase in RBR1 phosphorylation at Ser911, a western blot was performed on proteins extracted from root tips of WT and *CYCA3;4^OE^* homozygous line 12.4 using antibodies specifically targeting the phospho-Ser911 site and total RBR1. In both repeats, the ratio of S911-phosphorylated RBR1 on total RBR1 in the *CYCA3;4*-overexpressing background was twice that of the ratio in WT (Figures 9G and 9H).

## DISCUSSION

CCS52 proteins play an important role in restraining cell division through the stimulation of proteolytic turnover of proteins during the cell cycle. CCS52A2 in particular has a key function in preventing unscheduled stem cell divisions, as its deficiency results in a distorted stem cell niche, both in the root and shoot (Vanstraelen et al., 2009; Liu et al., 2012). Despite its developmental importance, the number of known or potential APC/C^CCS52A2^ targets is limited. Here, we identified through a suppressor screen the CYCA3;4 protein as a very likely proteolytic APC/C^CCS52A2^ target with an important role in controlling stem cell divisions. Knockout of *CYCA3;4* in the *ccs52a2-1* mutant background partially rescued the stem cell organization and root growth phenotypes, as well as leaf cell division defects. The data imply that the inability to control the CYCA3;4 protein level is one of the underlying reasons for the *ccs52a2-1* mutant phenotypes. Strikingly, introducing a *CYCA3;4* complementation construct in the *pkn2 ccs52a2-1* background or a translational reporter line within the *ccs52a2-1* mutant background predominantly resulted in an enhancement of the *ccs52a2-1* phenotype. We speculate that the origin of this enhanced phenotype might be the additional increase in CYCA3;4 abundance because of the introduction of one or more additional gene copies, again suggesting that that the level of CYCA3;4 abundance needs to be strictly controlled.

The evidence of CYCA3;4 being an APC/C^CCS52A2^ target is compelling. Not only does a mutation in *CYCA3;4* complement the *ccs52a2-1* phenotype, but co-overexpression with *CCS52A2* also suppresses the hyperproliferation phenotype of *CYCA3;4*-overexpressing plants. Moreover, the CYCA3;4 protein was previously found to co-immunoprecipitate with CCS52A2 (Fülöp et al., 2005). Additionally, we found that the *CYCA3;4_pro_:CYCA3;4-GUS* translational reporter protein predominantly accumulates in the distal root meristem following treatment with a proteasome inhibitor or when introduced within the *ccs52a2-1* mutant background, matching with the spatial accumulation pattern of CCS52A2. Finally, through immunostaining, the CYCA3;4-GUS protein could be detected on the chromosomes of metaphase and anaphase cells within the *ccs52a2-1* mutant background, whereas in WT cells the signal could only be detected in prophase nuclei. Next to strengthening the hypothesis that CYCA3;4 is an APC/C^CCS52A2^ target, these data also suggest that the APC/C^CCS52A2^ complex becomes active during mitosis, more precisely before metaphase.

Whereas knockout of *CYCA3;4* partially rescues the *ccs52a2-1* mutant phenotype, no outspoken phenotypes could be observed upon loss of CYCA3;4 activity in a WT background, suggesting redundancy with other cyclins. *CYCA3;4* is part of a gene family holding four members. *CYCA3;4* itself is part of a tandem duplication also containing *CYCA3;2* (AT1G47210) and *CYCA3;3* (AT1G47220), whereas *CYCA3;1* (AT5G43080) resides on a different chromosome. The different chromosomal localization of *CYCA3;1* and *CYCA3;4* suggests genetic diversification, which can be seen in the distinct spatial and temporal accumulation patterns of their respective proteins. Whereas CYCA3;1 predominantly accumulates in the proximal root meristem, CYCA3;4 can also be detected in the stem cell region. Its presence in the upper meristem marks CYCA3;1 as a putative target for APC/C^CCS52A1^ rather than APC/C^CCS52A2^, as CCS52A1 predominantly accumulates in the root at the beginning of the elongation zone, fitting with its role as a determinant of root meristem size (Vanstraelen et al., 2009). Correspondingly, a mutation in *cyca3;1* could not complement the *ccs52a2-1* phenotype, suggesting that CYCA3;1 is not an APC/C^CCS52A2^ substrate. Functional diversification between CYCA3;1 and CYCA3;4 is also supported by their differential temporal protein accumulation pattern, with CYCA3;1 and CYCA3;4 peaking during the S and G2/M phases, respectively. The *CYCA3;3* gene appears to be meiosis specific, as no transcript or protein could be detected in non-meiotic cells (Takahashi et al., 2010; Bulankova et al., 2013), leaving *CYCA3;2* as the most likely gene operating redundantly with *CYCA3;4*. Redundancy between *CYCA3;2* and *CYCA3;4* might explain why knockout of *CYCA3;4* did not completely rescue the *ccs52a2-1* root growth and meristem phenotype. Likewise, although *CYCA3;4* deficiency almost completely rescued the leaf cell division phenotype, only a mild rescue of the cell size and endoreplication phenotypes could be observed, suggesting the need to break down cyclins other than CYCA3;4 to enable normal cell growth and endocycle progression. Alternatively, the inability to degrade proteins other than cyclins might account for the residual phenotypes of the *cyca3;4 ccs52a2-1* double mutants. As such, previously ERF115, a transcription factor rate-limiting for QC stem cell division, and CSLD5, a cell wall biosynthesis enzyme involved in the production of the cell plate during division, were shown to be under proteolytic control of APC/C^CCS52A2^.

The need for controlled CYCA3;4 destruction is highlighted by the phenotypes triggered upon overexpression of the *CYCA3;4* gene, resulting in a small leaf phenotype. Remarkably, no lines with very high *CYCA3;4* transcript levels could be obtained and plants were prone to gene silencing, suggesting that strong overexpression might be counter-selected for, a situation also seen upon overexpression of the tobacco *CYCA3;2* gene (Yu et al., 2003). The small leaf phenotype of the *CYCA3;4*-overexpressing lines was mainly caused by a dramatic effect on cell size, being only partially offset by an increase in cell number. This makes the *CYCA3;4* overexpression phenotype different from that of the overexpression of other cyclins, such as *CYCD3;1*, in which the small cell phenotype is accompanied by a 20- to 30-fold increase in epidermal cell number (Dewitte et al., 2003), whereas for *CYCA3;4* only a maximum twofold increase in cell number was observed. Another major difference between *CYCD3;1*- and *CYCA3;4*-overexpressing lines is the lack of stomata in the latter. Therefore, it is possible that the cause of the reduction in cell size might not be the same. Whereas the small cell phenotype of the *CYCD3;1*-overexpressing plants might originate from an inability to enter cell differentiation, we speculate that within the CYCA3;4-overproducing lines this might be an indirect consequence of a reduction in carbon fixation due to the lack of stomata.

Next to the small leaf phenotype, *CYCA3;4*-overexpressing lines display an expression level-dependent root meristem phenotype. Whereas more highly overexpressing lines only display a short root meristem phenotype due to a reduction in the number of meristem cells, the lines with a lower overexpression also display an increased frequency of aberrant divisions, including unscheduled periclinal divisions. Combined with the effect on stomata, this suggests that CYCA3;4 might play an important role in the process of formative cell divisions, which might correspond to the need for its destruction by APC/C^CCS52A2^ to obtain a well-organized stem cell niche. Its targeted destruction during early prophase, the moment when the division plane orientation is determined through positioning of the preprophase band (Rasmussen and Bellinger, 2018; Facette et al., 2019), fits the idea that CYCA3;4 and CCS52A2 might play a role in the positioning of the division plane. However, the phenotype of the *CYCA3;4*-overexpressing plants does not completely mimic that of the *ccs52a2-1* knockout, again suggesting that the stabilization of targets other than CYCA3;4 might account for a big part of the disorganized stem cell niche phenotype of *ccs52a2-1* plants.

Strikingly, two of the phenotypes observed, being the stomata phenotype and the unscheduled stem cell divisions, are shared with plants silenced for the *RBR1* tumor suppressor gene (Wildwater et al., 2005; Borghi et al., 2010; Cruz-Ramírez et al., 2012; Matos et al., 2014). Reciprocally, hypomorphic *CDKA;1* mutants have been described to display delayed formative divisions in both the root and shoot, and this could be suppressed by a mutation in the *RBR1* gene (Weimer et al., 2012). Because it is anticipated that phosphorylation by CDK proteins inhibits RBR1 activity (Harashima and Sugimoto, 2016), these data suggest that RBR1 inactivation induces formative divisions. Through our phosphoproteomics analysis of *CYCA3;4* overexpression plants, an enrichment for two phospho-sites within the RBR1 protein (T406 and S911) could be found. Both identified sites are part of the minimal CDK phosphorylation site [S/T]P and are highly conserved, corresponding to respectively T373 and S807 within the human RB protein, for which their phosphorylation has been demonstrated to reduce RBR’s inhibitory binding of E2F transcription factors (Ren and Rollins, 2004; Burke et al., 2010; Burke et al., 2012; Burke et al., 2014). Assuming that the phosphorylation of RBR1 triggers an identical effect, it might be speculated that CYCA3;4 in complex with CDKA;1 might regulate stem cell identity or polarity of cell divisions through inhibition of RBR1, and that this activity is restrained through APC/C^CCS52A2^ activity. Furthermore, as only a limited number of proteins were found to display increased phosphorylation upon *CYCA3;4* overexpression, it indicates that RBR1 might be the main target of CYCA3;4. However, we currently do not have biochemical evidence that RBR1 is a direct target of a CYCA3;4-CDKA;1 pair, as through interaction experiments we failed to detect direct binding between RBR1 and CYCA3;4, fitting with the absence of an LxCxE RBR1 interaction motif in CYCA3;4. Therefore, it currently cannot be excluded that the increase in RBR1 phosphorylation might be an indirect consequence of the strong phenotypic effects of *CYCA3;4* overexpression. Interestingly, expression of the *CCS52A* genes is under direct negative control of the RBR1 protein (Magyar et al., 2012), leading to the possibility that CYCA3;4 might be responsible for triggering its own APC/C^CCS52A2^-mediated breakdown through the phosphorylation and inactivation of RBR1.

## METHODS

### Plant Medium and Growth Conditions

Seeds of *Arabidopsis thaliana* were sterilized by incubating in 70% ethanol for 10-15 min and subsequent washing with 100% ethanol, after which they were left to dry in sterile conditions. Plants were grown in vitro under long-day conditions (16-h light/8-h dark) at 21°C on half-strength Murashige and Skoog (MS) medium (2.151 g/L), 10 g/L sucrose, and 0.5 g/L 2-(N-morpholino) ethanesulfonic acid (MES), adjusted to pH 5.7 with 1 M KOH and 10 g/L agar. For analysis of root or shoot phenotypes, plants were grown vertically or horizontally, respectively. The drug treatments described were performed using the following conditions: MG132 – 100 μM for 24 h; hydroxyurea – 1 mM for 24 h; bleomycin – 0.9 μg/mL (0.636 μM) for 24 h; oryzalin – 0.3 μM for 24 h.

### Constructs and Lines

The mutant lines *ccs52a1-1, ccs52a2-1* and *cyca3;1-1* have been described previously (Vanstraelen et al., 2009; Takahashi et al., 2010), whereas *cyca3;4-2* (SALK_204206) and *cyca3;4-3* (SALK_061456) were obtained from the Salk Institute T-DNA Express (Alonso et al., 2003) database. The *pkn2 ccs52a2-1* double mutant was obtained through EMS mutagenesis of *ccs52a2-1* mutant seeds (see below). The *CYCA3;4* complementation construct was created by cloning a fragment containing the *CYCA3;4* promoter and gene sequence from Col-0 into the pDONR211 vector and recombining it into the pFAST-R01 vector. The *CYCA3;4^OE^* construct was created by cloning the *CYCA3;4* open reading frame (ORF) from Col-0 into pDONR221 and subsequently recombining it behind the strong CaMV 35S promoter in the pB7WG2 vector. The *CCS52A2^OE^* line was kindly donated by Dr. Eva Kondorosi (Baloban et al., 2013). The *CYCA3;1_pro_:CYCA3;1-GUS* and *CYCA3;4_pro_:CYCA3;4-GUS* translational reporter lines were kindly donated by Dr. Karl Riha (Bulankova et al., 2013). The *CCS52A1_pro_:CCS52A1-GUS* and *CCS52A2_pro_:CCS52A2-GUS* translational reporter constructs were created by cloning a fragment consisting of the 2000-bp sequence upstream of the start codon followed by the complete gene including introns but without stop codon into the pDONR^™^-P4-P1r entry vector and cloning it in front of the GUS reporter by recombining it with pEN-L1-SI*-L2 into the pK7m24GW-FAST vector. All primer sequences used for cloning and genotyping are listed in Supplemental Table 3.

All vector-based cloning was performed using the Gateway system (Invitrogen). All constructs were transformed into the Col-0 background using the floral dipping technique, except the *CYCA3;4* complementation construct, which was transformed into *pkn2 ccs52a2-1*. Successful transformants were selected using Kanamycin or Basta, or using fluorescent microscopy in case of FAST constructs. Double mutants were made by crossing and confirmed through genotyping with PCR and/or sequencing.

### Plant Growth Phenotyping

Root growth and length were determined by marking the position of the root tip each day from 3 to 9 DAS, scanning the plates at 9 DAS and measuring using the ImageJ software package. Root meristem analysis was performed with the ImageJ software package using images of the root tip obtained with confocal microscopy, the distance between the QC and the end of the division zone was measured to determine the root meristem length, and the number of cortical cells within the division zone was counted to determine the cortical cell number.

For rosette size, pictures were taken at 21 DAS using a digital camera fixed in position, after which the images were made binary (black and white) and the projected rosette size was measured using the wand tool in ImageJ. For analysis of leaf parameters, the first leaf pairs were harvested at 21 DAS and cleared overnight using 100% ethanol. Next, leaves were mounted on a slide with lactic acid. The total leaf area was determined from images taken with a digital camera mounted on a Stemi SV11 microscope (Zeiss) using the ImageJ software. Using a darkfield microscope (Leica) with a drawing-tube attached, an area of the leaf was drawn containing the outlines of at least 30 cells of the abaxial epidermis located between 25 to 75% of the distance between the tip and the base of the leaf, halfway between the midrib and the leaf margin. After measuring the total drawn area (using the wand tool of ImageJ) and counting the number of pavement cells and stomata drawn, the average cell size, total number of cells per leaf and the stomatal index (number of stomata divided by total number of epidermal cells) were calculated.

Statistical analyses were done with the SAS Enterprise Guide 7 software using the ANOVA or Mixed Model procedure and Tukey- or Dunnett-correction for multiple testing.

### Flow Cytometry

For flow cytometry analysis, leaf material was chopped in 200 μL nuclei extraction buffer, after which 800 μL staining buffer was added (Cystain UV Precise P, Partec). The mix was filtered through a 30-μm green CellTrics filter (Sysmex – Partec) and analyzed by the Cyflow MB flow cytometer (Partec). The Cyflogic software was used for ploidy measurements.

### Confocal Microscopy

For visualization of root apical meristems, vertically grown plants were mounted in a 10-μM propidium iodide (PI) solution (Sigma) to stain the cell walls and imaged using an LSM 5 Exciter (Zeiss) confocal microscope. For PI and GFP or YFP excitation, the 543 line of a HeNe laser and the 488 line of an Argon laser were used, respectively. Laser light passed through an HFT 405/488/543/633 beamsplitter before reaching the sample, and emitted light from the sample first passed through an NFT 545 beamsplitter, after which it passed through a 650-nm long pass filter for PI detection, and through a 505- to 530-nm band pass filter for detection of GFP or YFP. PI and GFP or YFP were detected simultaneously with the line scanning mode of the microscope.

### GUS Staining

Plants were grown for the indicated time and fixated in an ice-cold 80% acetone solution for 3 h. Samples were washed three times with phosphate buffer (14 mM NaH_2_PO_4_ and 36 mM Na_2_HPO_4_) before being incubated in staining buffer (0.5 mg/mL 5-bromo-4-chloro-3-indolyl-β-D-glucuronic acid, 0.165 mg/mL potassium ferricyanide, 0.211 mg/mL potassium ferrocyanide, 0,585 mg/mL EDTA pH8, and 0,1% (v/v) Triton-X100, dissolved in phosphate buffer) at 37°C between 30 min and 16 h until sufficient staining was observed.

### EMS Mutagenesis

Roughly 14,000 *ccs52a2-1* seeds were subjected to EMS mutagenesis. The seeds were first hydrated with H_2_O for 8 h on a rotating wheel before being mutagenized with a 0.25% v/v solution of EMS for another 12 h. After treatment, seeds were washed twice with 15 mL 0.1 M sodium thiosulfate (Na_2_S_2_O_3_) for 15 min to stop the reaction and subsequently twice with H_2_O for 30 min. After that, seeds were left to dry on Whatman paper. Fifty-six pools of approximately 250 M_1_ seeds were mixed together with fine sand in Eppendorf tubes and sown in big pots with standard soil. After selfing, M_2_ seeds were harvested separately for every pool. Seeds were sterilized and sown on vertical plates to score for the reversion of the *ccs52a2-1* root growth phenotype. Plants with longer roots were subsequently selected and transferred to soil for self-fertilization. The recovery phenotype was then reconfirmed in the next generation (M_3_).

### Mapping of the Revertant Mutation

Segregating F2 progeny resulting from a cross between *pkn2 ccs52a2-1* and the *ccs52a2-1* parental line used for EMS mutagenesis was used as a mapping population. Approximately 250 plants showing the long root phenotype of the revertant were selected at 5 DAS and pooled for DNA extraction using the DNeasy Plant Mini Kit (Qiagen) according to the manufacturer’s instructions. DNA was extracted additionally from 200 plants of the original *ccs52a2-1* parental line. Illumina True-Seq libraries were generated from the extracted DNA according to the manufacturer’s protocol and sequenced on an Illumina HiSeq 100-bp paired-end run. The SHORE pipeline (Ossowski et al., 2008) was used for the alignment of sequences of both *pkn2 ccs52a2-1* and *ccs52a2-1* to the reference genome (Col-0; TAIR10). Using the SHOREmap pipeline (Sun and Schneeberger, 2015), an interval of increased mutant SNP alleles was identified and subsequently annotated. Filtering was performed within the interval for *de novo* EMS-specific SNPs with a concordance above 0.8 and intergenic or intronic mutations were removed to reveal the potential revertant mutations.

### RT-PCR and qRT-PCR

RNA was isolated with the RNeasy Mini kit (Qiagen) and was treated on-column with the RQ1 RNase-Free DNase (Promega). Synthesis of cDNA was done with the iScript cDNA Synthesis Kit (Bio-Rad). For visualization of the *CYCA3;4* splice variants created by the EMS mutation, cDNA from *pkn2 ccs52a2-1* and Col-0 was separated on a 1% agarose gel containing SYBRSafe (Invitrogen). For determination of relative expression levels, the LightCycler 480 Real-Time SYBR Green PCR System (Roche) was used. Three reference genes were used for normalization: *EMB2386, PAC1* and *RPS26E*. All primer sequences used for qRT-PCR are listed in Supplemental Table 3.

### Immunostaining Experiment

Plants were grown vertically on full strength MS medium (supplemented with 20 g/L sucrose, 0.1 g/L myo-inositol, 0.5 g/L MES, 10 g/L thiamine hydrochloride, 5 g/L pyridoxine, 5 g/L nicotinic acid, pH adjusted to 5.7 with 1 M KOH, and 10 g/L plant agar) for 9 days. Root tips were fixed for 45 min in 4% paraformaldehyde in a solution of 1X PME (50 mM Pipes pH 6.9, 5 mM MgSO_4_, 1 mM EGTA) and then washed three times for 5 min in 1X PME. Root apices were dissected on a glass slide and digested in a drop of enzyme mix (1% w/v cellulase, 0.5% w/v cytohelicase, 1% w/v pectolyase in PME) for 1 h at 37°C. After three washes with PME, root apices were squashed gently between the slide and a coverslip, and frozen in liquid nitrogen. Afterwards, the coverslip was removed and the slides were left to dry for 1 h at room temperature.

For immunostaining, each slide was incubated overnight at 4°C with 100 μL of rabbit polyclonal anti-β-glucuronidase antibody (N-Terminal, 5420, Molecular Probes, Invitrogen) diluted 1:200 in fresh blocking buffer (3% BSA in 1X PBS). Slides were washed three times for 5 min in 1X PBS solution and then incubated for 3 h at room temperature in 100 μL blocking buffer containing Alexa 488-conjugated goat anti-rabbit secondary antibody (Molecular Probes, Invitrogen), diluted 1:200 in fresh blocking buffer. Finally, DNA was counterstained with 2 μg/mL DAPI for 30 min, after which slides were washed in 1X PBS and mounted in mounting medium. The microscope settings and exposure times were kept constant for each respective channel.

### Phosphoproteomics

#### Protein Extraction and Phosphopeptide Enrichment

Total protein extraction was conducted on three biological replicates of approximately 50 pooled 14-DAS-old whole *CYCA3;4^OE^* 11.2 x Col-0 and Col-0 x Col-0 F1 seedlings, as previously described (Vu et al., 2017). Phosphopeptides were enriched as previously described with minor modifications (Vu et al., 2017). A total of 100 μL of the re-suspended peptides were incubated with 3 mg MagReSyn Ti-IMAC microspheres for 20 min at room temperature. The microspheres were washed once with wash solvent 1 (80% acetonitrile, 1% TFA, 200 mM NaCl) and twice with wash solvent 2 (80% acetonitrile, 1% TFA). The bound phosphopeptides were eluted with three volumes (80 μL) of an elution solution (40% ACN, 1% NH4OH, 2% formic acid), immediately followed by acidification to pH≤ 3 using 100% formic acid. Prior to MS analysis, the samples were vacuum dried and re-dissolved in 50 μL of 2% (v/v) acetonitrile and 0.1% (v/v) TFA, of which 10 μL was injected for LC-MS/MS analysis.

#### LC-MS/MS Analysis

Each sample was analyzed via LC-MS/MS on an Ultimate 3000 RSLC nano LC (Thermo Fisher Scientific) in-line connected to a Q Exactive mass spectrometer (Thermo Fisher Scientific). The peptides were first loaded on a trapping column (made in-house, 100-μm internal diameter (I.D.) × 20 mm, 5-μm beads C18 Reprosil-HD, Dr. Maisch, Ammerbuch-Entringen, Germany). After flushing the trapping column, peptides were loaded in solvent A (0.1% formic acid in water) on a reverse-phase column (made in-house, 75-μm I.D. x 250 mm, 1.9-μm Reprosil-Pur-basic-C18-HD beads, Dr. Maisch, packed in the needle) and eluted by an increase in solvent B (0.1% formic acid in acetonitrile) using a linear gradient from 2% solvent B to 55% solvent B in 180 min, followed by a washing step with 99% solvent B, all at a constant flow rate of 300 nL/min. The mass spectrometer was operated in data-dependent, positive ionization mode, automatically switching between MS and MS/MS acquisition for the five most abundant peaks in a given MS spectrum. The source voltage was set at 4.1 kV and the capillary temperature at 275°C. One MS1 scan (m/z 400-2,000, AGC target 3 × 106 ions, maximum ion injection time 80 ms), acquired at a resolution of 70,000 (at 200 m/z), was followed by up to five tandem MS scans (resolution 17,500 at 200 m/z) of the most intense ions fulfilling predefined selection criteria (AGC target 5 × 104 ions, maximum ion injection time 80 ms, isolation window 2 Da, fixed first mass 140 m/z, spectrum data type: centroid, under-fill ratio 2%, intensity threshold 1.3xE4, exclusion of unassigned, 1, 5-8, >8 positively charged precursors, peptide match preferred, exclude isotopes on, dynamic exclusion time 12 s). The HCD collision energy was set to 25% Normalized Collision Energy and the polydimethylcyclosiloxane background ion at 445.120025 Da was used for internal calibration (lock mass).

#### Database Searching

MS/MS spectra were searched against the Arabidopsis database downloaded from TAIR with the MaxQuant software (version 1.5.4.1), a program package allowing MS1-based label-free quantification acquired from Orbitrap instruments (Cox and Mann, 2008; Cox et al., 2014). Next, the ‘Phospho(STY).txt’ output file generated by the MaxQuant search was loaded into the Perseus data analysis software (version 1.5.5.3) available in the MaxQuant package. Proteins that were quantified in at least two out of three replicates from each crossed line were retained. Log2 phosphopeptide intensities were centered by subtracting the median. A two-sample test with a P-value cut-off of P < 0.01 was carried out to test for differences between the crossed lines. Additionally, phosphopeptides with three valid values in each crossed line and none in the other were also retained and designated “unique” for that condition. All MS proteomics data in this study have been deposited to the ProteomeXchange Consortium via the PRIDE partner repository (Vizcaino et al., 2016) with the data set identifier PXD017905.

### Western Blot

For western blot analysis, seeds were sown on nylon meshes (Prosep) on half strength MS medium supplemented with 2% sucrose. Approximately 5 mm root tips from one-week-old plants were harvested for protein extraction. Fifty micrograms total protein extracts were separated by means of SDS-PAGE and subsequently subjected to western blotting. Protein blots were hybridized with anti-RBR1 (Agrisera; #AS11 1627) and anti-Phospho-RB (Ser807/811) (Cell Signaling Technology; #8516T) antibodies according to the manufacturer’s description. Protein levels were quantified from two biological repeats, using three different exposures obtained from each repeat, using the ImageJ analysis software.

### Accession Numbers

Sequence data from this article can be found in the Arabidopsis Genome Initiative or GenBank/EMBL databases under the following accession numbers: *CCS52A1* (AT4G22910), *CCS52A2* (AT4G11920), *CYCA3;1* (AT5G43080), *CYCA3;4* (AT1G47230), WOX5 (AT3G11260), SPCH (AT5G53210), SDD1 (AT1G04110), MUTE (AT3G06120), FAMA (AT3G24140), *RBR1* (AT3G12280), EMB2386 (AT1G02780), PAC1 (AT3G22110) and RPS26E (AT3G56340).

## Supplemental Data

The following materials are available in the online version of this article.

**Supplemental Data Set 1.** List of identified phosphosites from phosphoprofiling of Col-0 x Col-0 (WT) and *CYCA3;4^OE^* 11.2 x Col-0 (OE) seedlings

**Supplemental Data Set 2.** List of phosphosites significantly deregulated (Students’ t-test p < 0.01) in Col-0 x Col-0 (WT) versus *CYCA3;4^OE^* 11.2 x Col-0 (OE) seedlings

**Supplemental Data Set 3.** List of “unique” deregulated phosphosites from phosphoprofiling of Col-0 x Col-0 (WT) versus *CYCA3;4^OE^* 11.2 x Col-0 (OE) seedlings

## ACKNOWLEDGMENTS

The authors thank Annick Bleys for help in preparing the manuscript. This work was supported by grants of the Research Foundation Flanders (G023616N and G007218N). A.W. is indebted to the Agency for Innovation by Science and Technology in Flanders for a predoctoral Fellowship. J.H. is indebted to the Research Foundation Flanders for a postdoctoral fellowship.

## AUTHOR CONTRIBUTIONS

A.W., J.H., I.D.S. and L.D.V. conceived and designed the research. A.W., J.H., T.E., I.A., J.A.P.G., T.Z, L.L., H.V.d.D., I.V., and B.v.d.C. performed the experiments. A.W., J.H., T.E., J.A.P.G, I.D.S. and L.D.V. analyzed the data. A.W. and L.D.V. wrote the article. All authors read, revised, and approved the article.

---

**Supplemental Figure 1.**
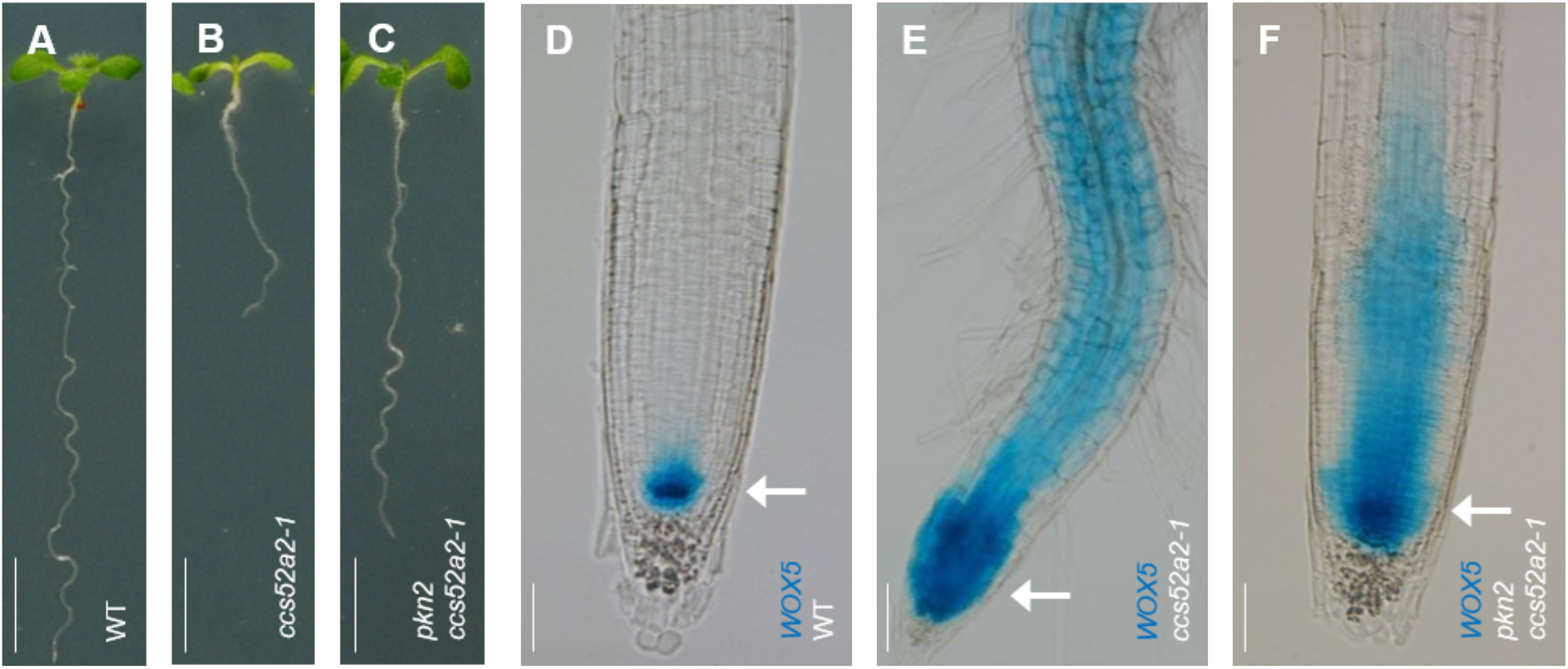
Additional Characteristics of the *ccs52a2-1* and *pkn2 ccs52a2-1* Mutants (Supports Figure 1). **(A-C)** Representative seedlings of WT **(A)**, *ccs52a2-1* **(B)** and *pkn2 ccs52a2-1* **(C)** at 9 DAS, showing recovery of the *ccs52a2-1* root growth phenotype in the double mutant. Scale bars represent 5 mm. **(D-F)** Visualization of the *WOX5_pro_:GFP-GUS* construct with histochemical GUS staining in the primary root of WT **(D)**, *ccs52a2-1* **(E)** and *pkn2 ccs52a2-1* **(F)** at 5 DAS, showing a strong ectopic *WOX5* expression in the *ccs52a2-1* background, which is much less severe in the double mutant. Arrows indicate the position of the QC. Scale bars represent 50 μm.

**Supplemental Figure 2.**
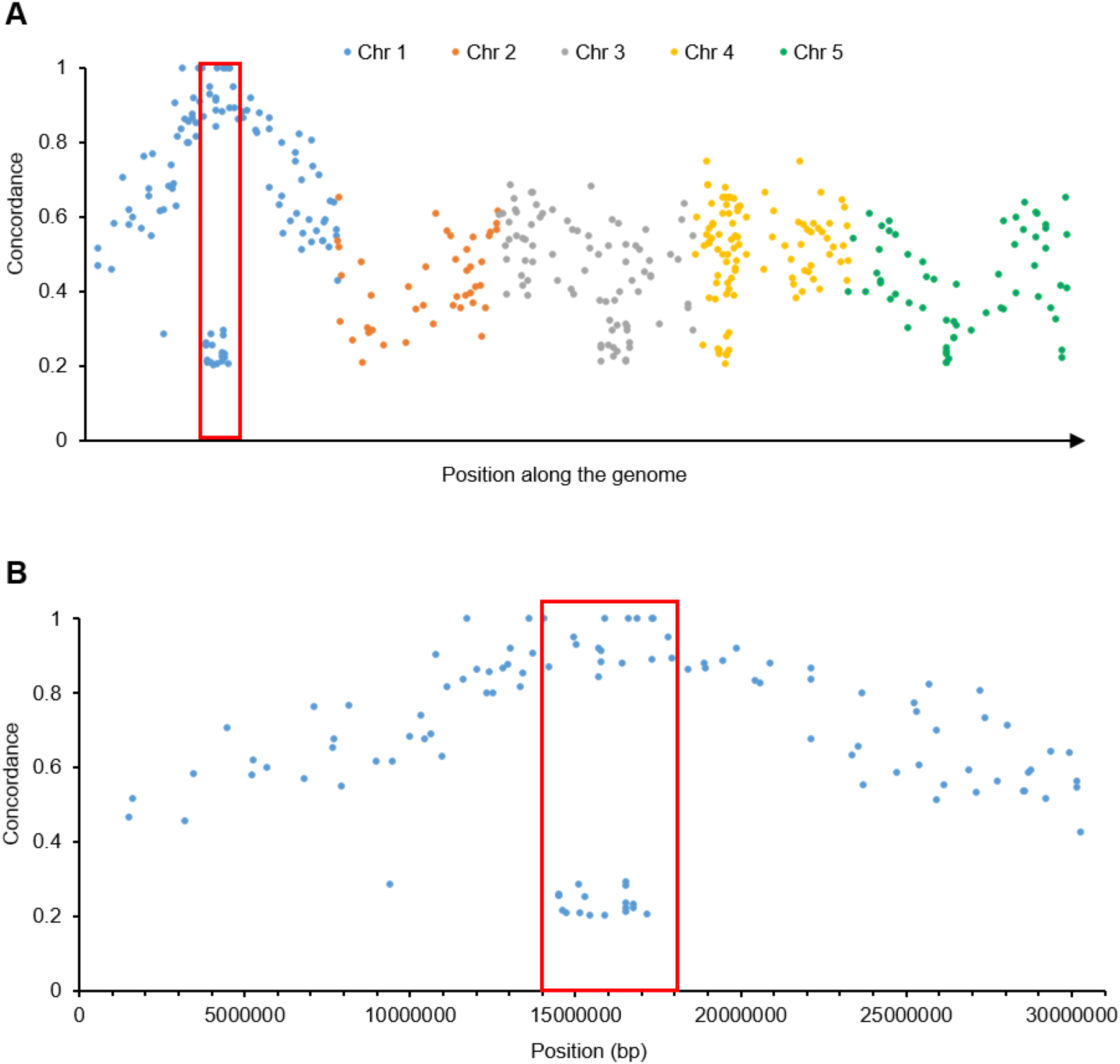
Detail of the Allele Frequency of EMS-Specific Mutations in *pkn2 ccs52a2-1* (Supports Figure 2). Allele frequency of mutant over WT alleles of newly identified EMS-specific mutations in function of their position on the whole *pkn2 ccs52a2-1* genome **(A)** and in detail on chromosome 1 **(B)**. The interval selected for a detailed annotated analysis is indicated by a red box. Identified mutations were filtered for uniqueness (i.e. not present in *ccs52a2-1*), EMS specificity (*i.e*. only G→A or C→T), quality (Q > 20) and coverage (reads > 10).

**Supplemental Figure 3.**
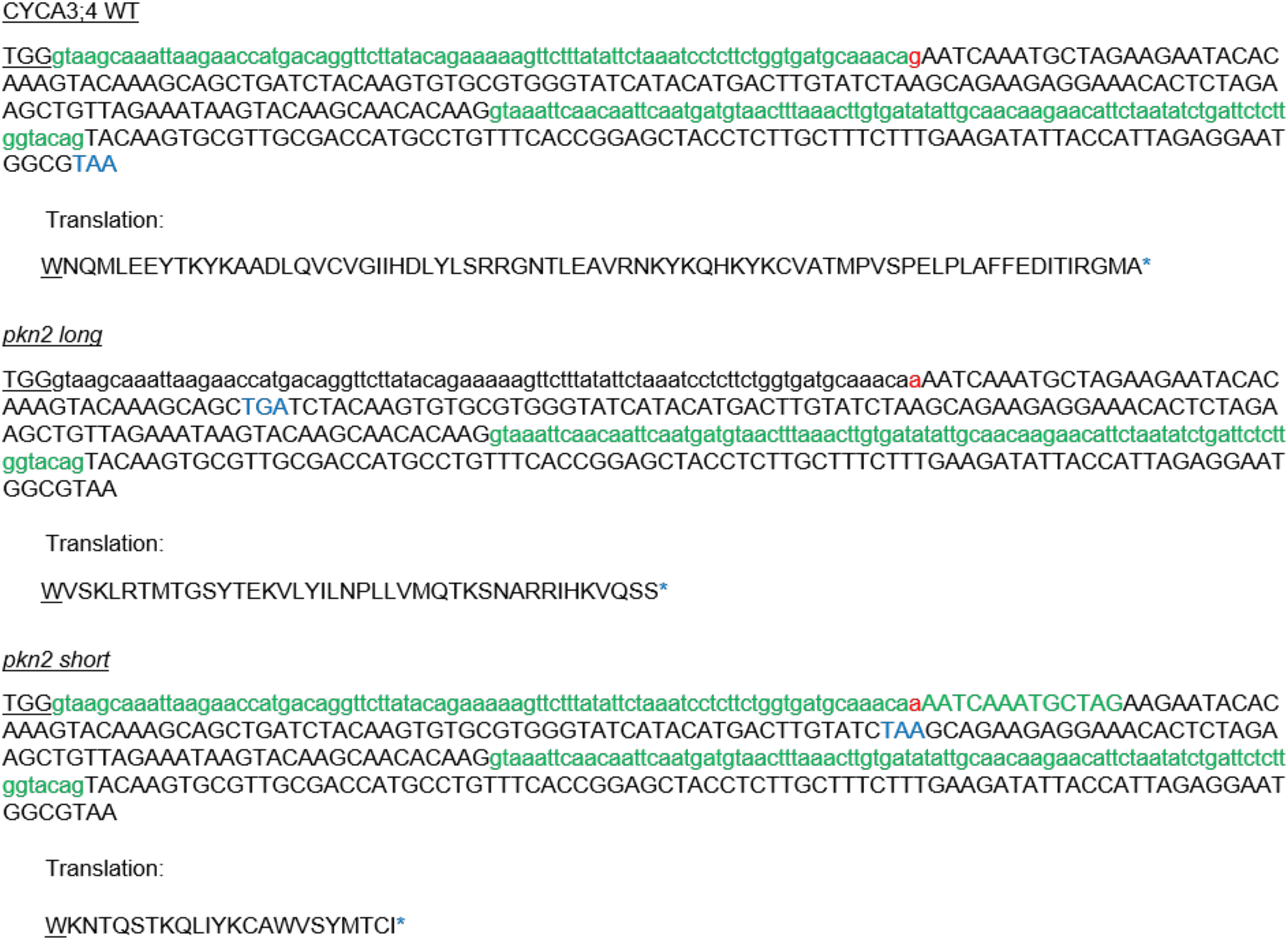
The *pkn2* EMS Mutation in *CYCA3;4* Causes Two Different Splice Variants to Be Expressed (Supports Figure 2). Detail of *CYCA3;4* from the last codon of exon 5 (underlined) to the end of the gene, showing the effect of the mutation at the splice acceptor site (marked in red) on mRNA and protein level. The introns (as spliced out in WT) are indicated with small letters; the stop codons are marked in blue. Spliced-out sequences are marked in green.

**Supplemental Figure 4.**
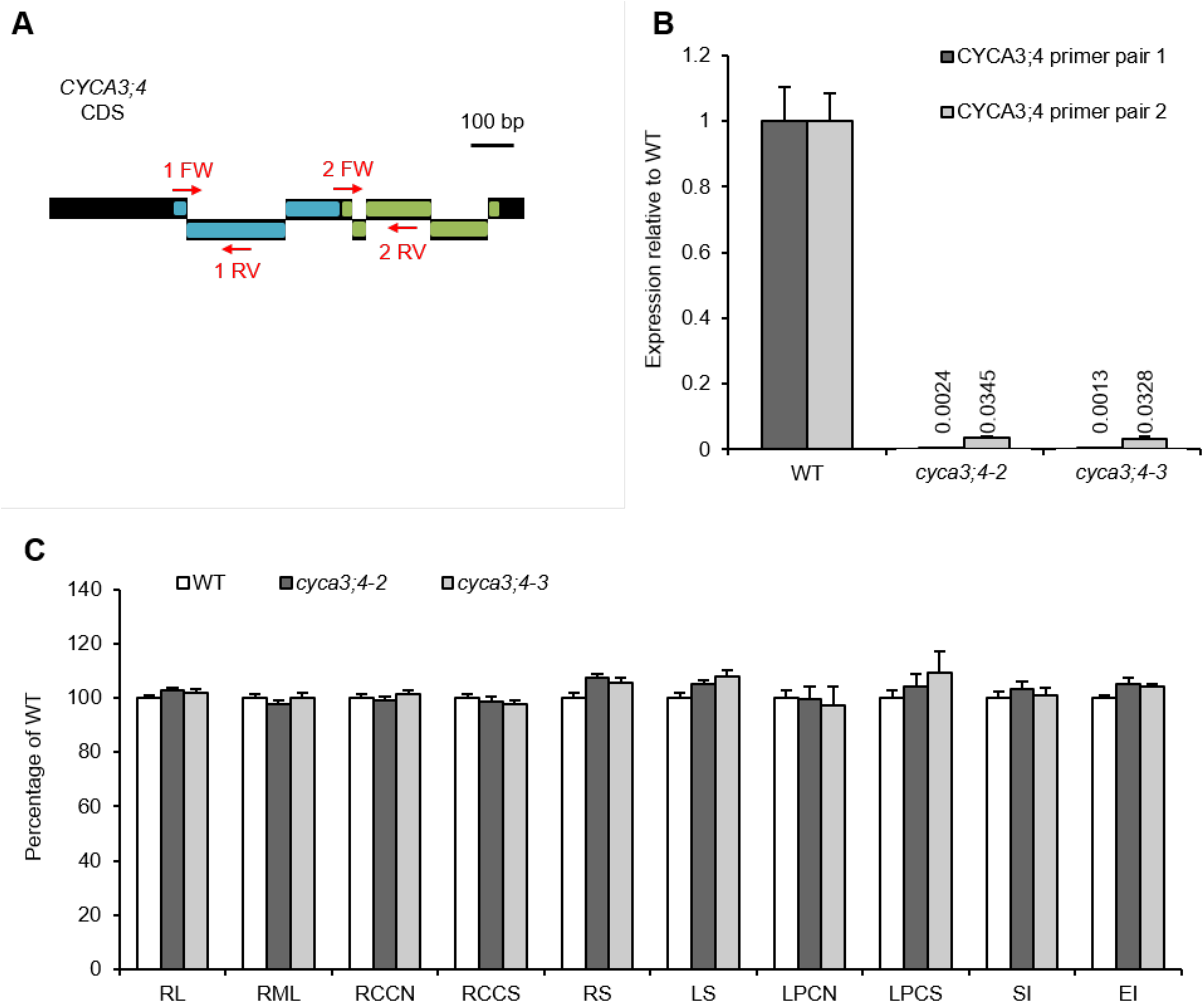
ANALYSIS OF *CYCA3;4* T-DNA INSERTION MUTANTS (SUPPORTS FIGURE 2). **(A)** The binding sites of the two primer pairs (red arrows) used for qRT-PCR on the *CYCA3;4* coding sequence. Primer pair 1 binds on the exon1-exon2 border and inside exon 2, while primer pair 2 binds on the exon3-exon4 border and exon 5. The predicted N- and C-terminal cyclin boxes are colored blue and green, respectively. **(B)** Expression levels of *CYCA3;4* relative to WT in the different T-DNA insertion lines, as measured by qRT-PCR using two different primer pairs for *CYCA3;4*. Because expression levels are very low, the exact values are indicated above the bar. Error bars represent standard error (n = 6). **(C)** Phenotypical analysis of *CYCA3;4* mutant lines *cyca3:4-2* and *cyca3;4-3* in the root (at 9 DAS) and shoot (at 21 DAS): primary root length (RL), root meristem length (RML), root cortical cell number (RCCL), projected rosette size (RS), leaf size of the first leaf pair (LS), leaf pavement cell number (LPCN), leaf pavement cell size (LPCS), stomatal index (SI) and endoreplication index (EI). Error bars represent standard error (n ≥ 5).

**Supplemental Figure 5.**
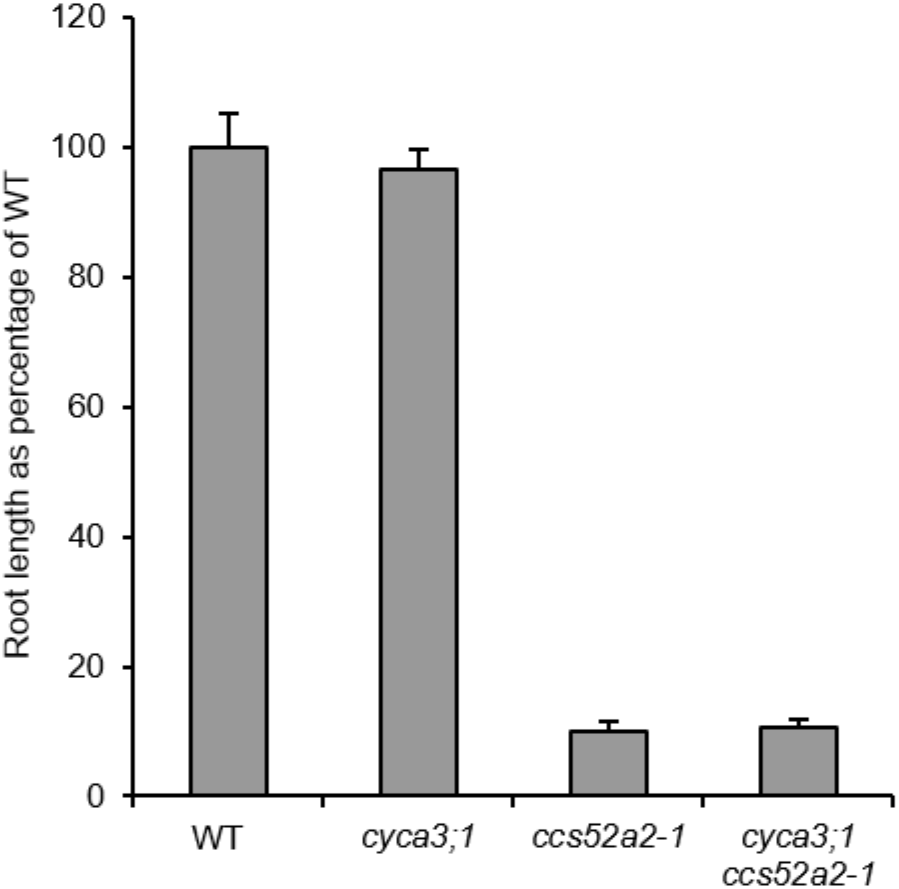
Lack of *CYCA3;1* Does Not Recover the *ccs52a2-1* Root Length Phenotype (Supports Figure 2). Root length at 9 DAS of WT, *cyca3;1, ccs52a2-1* and the double mutant *cyca3;1 ccs52a2-1*. Error bars represent standard error (n ≥ 12). Plants were genotyped and measured in the segregating F2 generation.

**Supplemental Figure 6.**
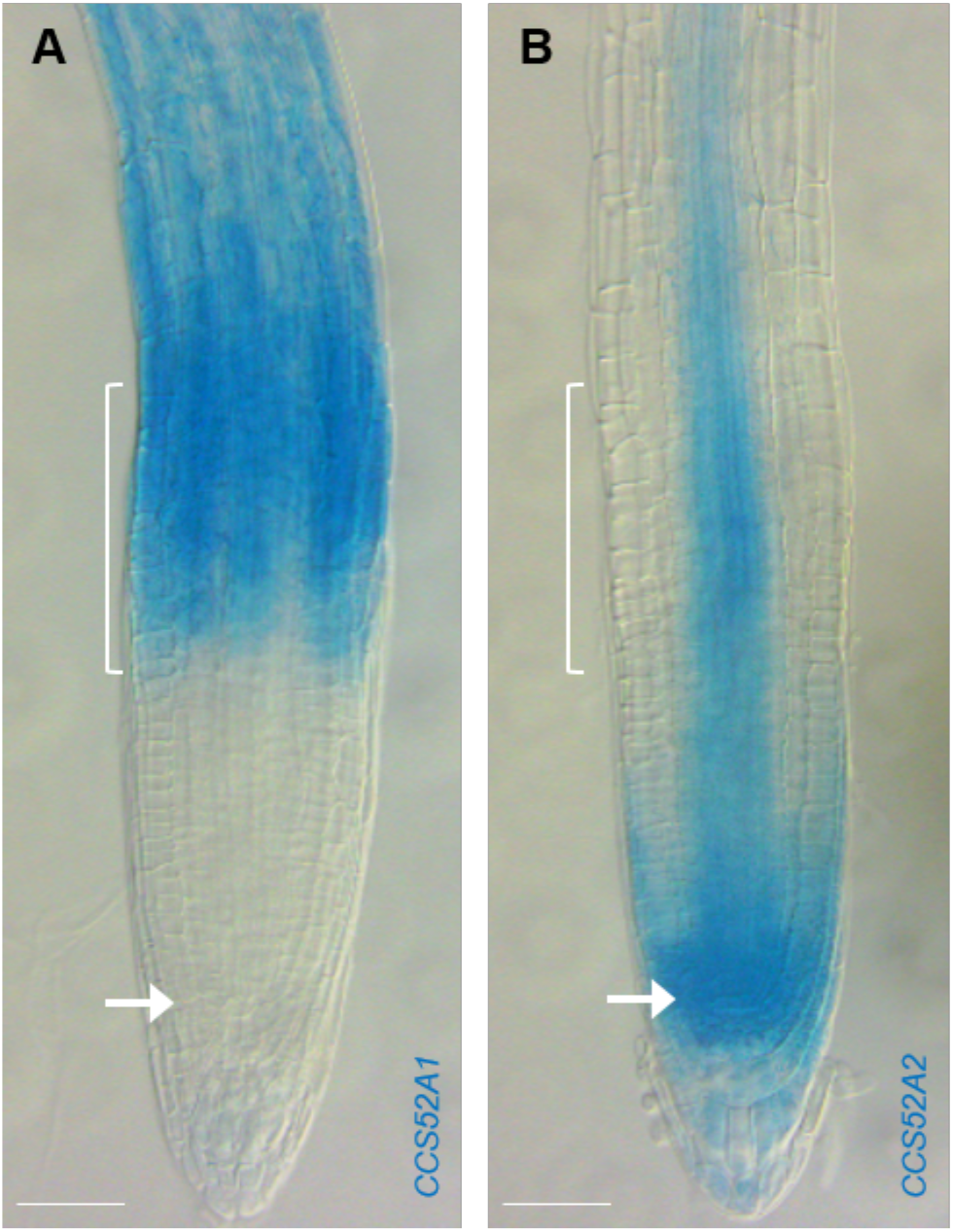
Root Tip Expression Pattern of the CCS52A Proteins (Supports Figure 3). Histochemical GUS staining using the translational reporter lines *CCS52A1_pro_:CCS52A1-GUS* **(A)** and *CCS52A2_pro_:CCS52A2-GUS* **(B)**, showing their expression at 5 DAS. The transition zone and the QC are indicated by brackets and arrows, respectively. Scale bars represent 50 μm.

**Supplemental Figure 7.**
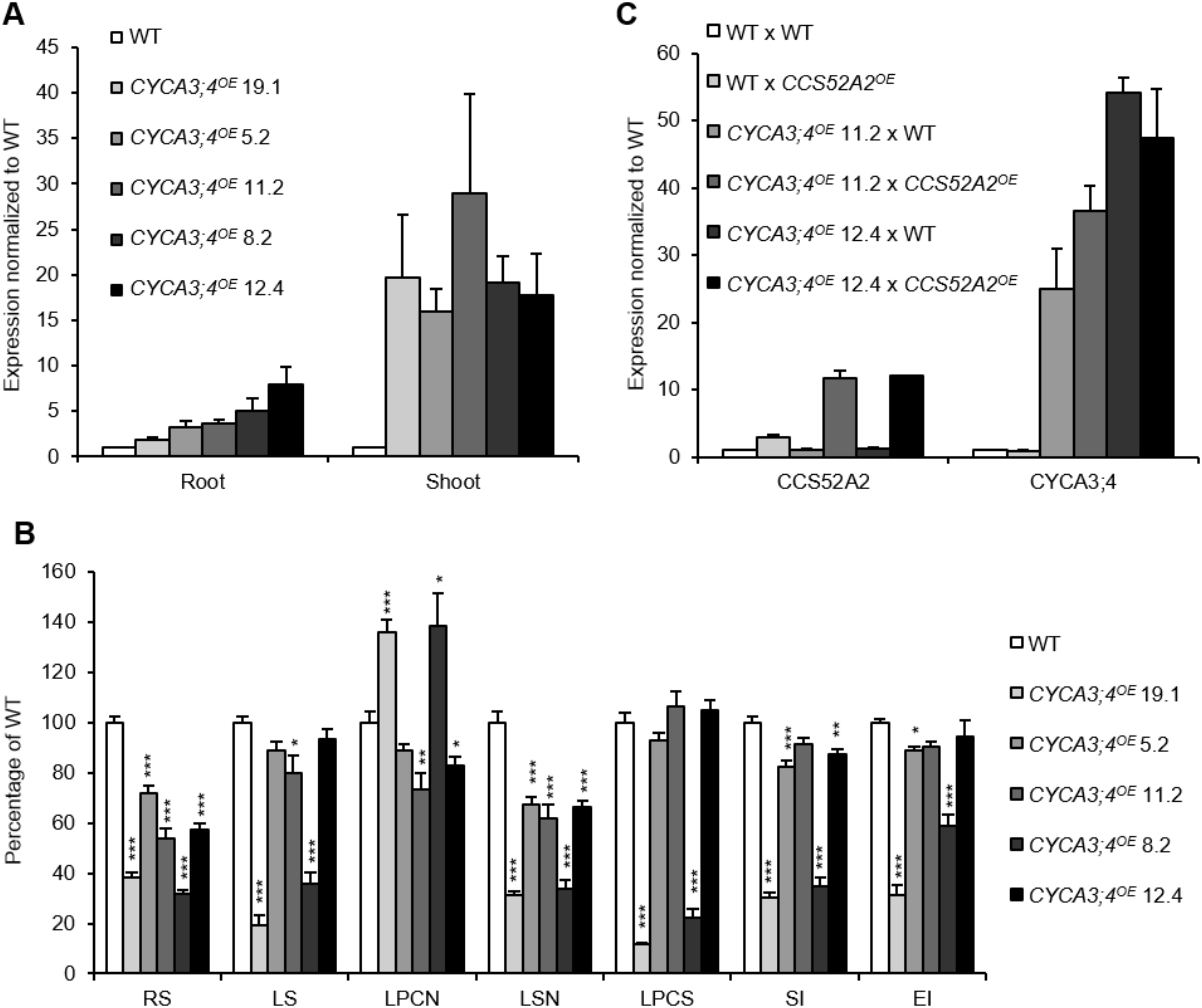
Expression Levels and Phenotypes in Different *CYCA3;4^OE^* Lines (Supports Figures 7 and 8). **(A)** *CYCA3;4* expression levels in homozygous T3 generation plants of the five different *CYCA3;4^OE^* lines, as measured by qRT-PCR on root tips at 9 DAS or whole shoots at 10 DAS. Error bars represent standard error (n = 2 or 3). **(B)** Shoot phenotypes at 21 DAS of homozygous T3 generation plants of the five different *CYCA3;4^OE^* lines: RS, rosette size; LS, leaf size; LPCN, leaf pavement cell number; LSN, leaf stomatal number; LPCS, leaf pavement cell size; SI, stomatal index; and EI, endoreplication index. Error bars represent standard error (n ≥ 12). Statistical significance was evaluated by a mixed model analysis, using Tukey correction for multiple testing: * p < 0.05, ** p < 0.01, *** p < 0.001. **(C)** *CCS52A2* and *CYCA3;4* expression levels in first-generation hemizygous progeny of crosses between two *CYCA3;4^OE^* lines and WT or *CCS52A2^OE^*, as measured by qRT-PCR in the first leaf pair at 21 DAS. Error bars represent standard error (n = 2 or 3).

**Supplemental Table 1.**
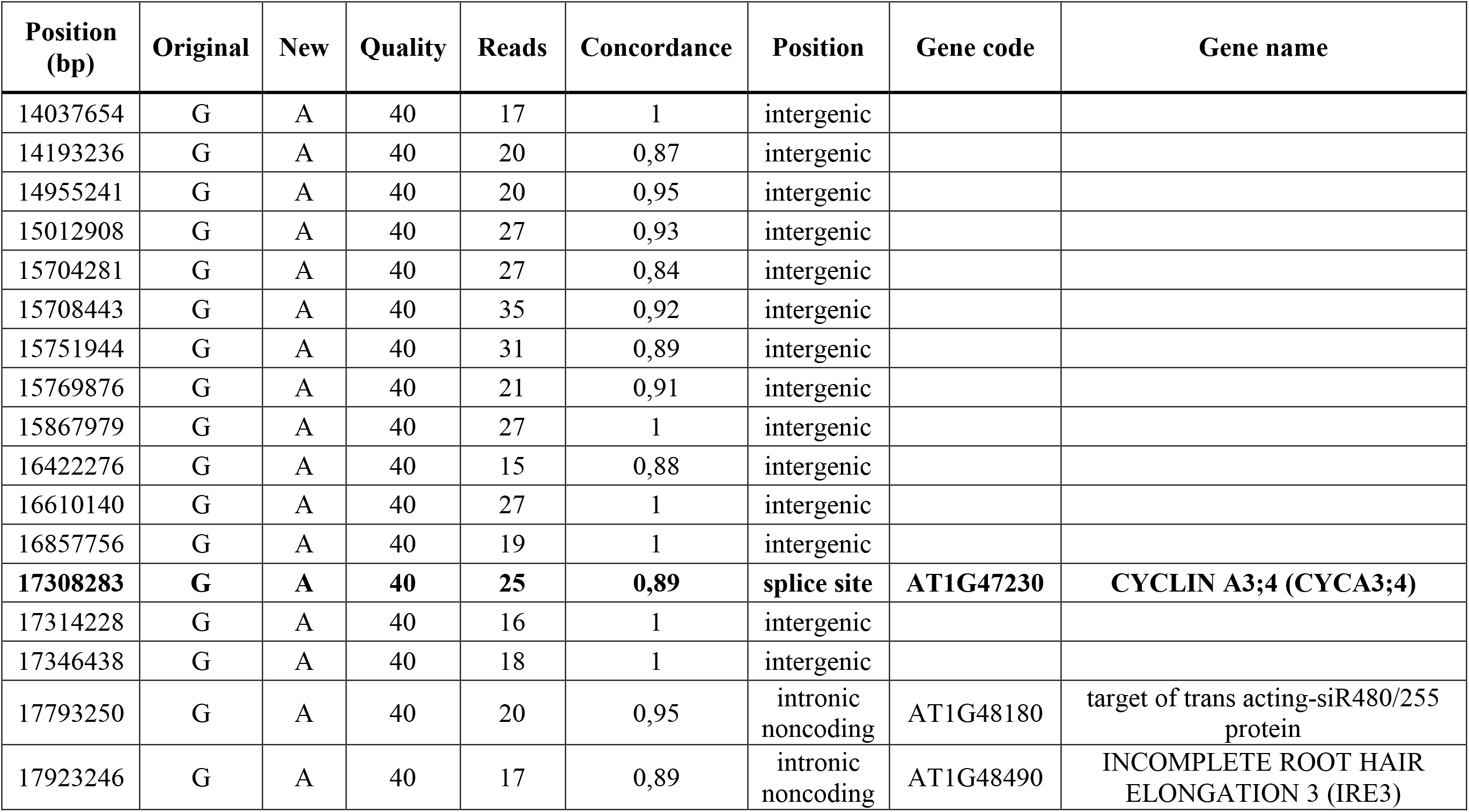
Detailed Annotation of the SNPs Found for *pkn2 ccs52a2-1* in the Interval Selected on Chromosome 1 from 14 Mbp to 18 Mbp. SNPs were filtered for uniqueness (i.e. not present in *ccs52a2-1*), EMS specificity (*i.e*. only G→A or C→T), quality (Q > 20), coverage (reads > 10) and concordance (C > 0,80). TAIR 10 version of the Arabidopsis Reference Genome.

| Position (bp) | Original | New | Quality | Reads | Concordance | Position | Gene code | Gene name |
| --- | --- | --- | --- | --- | --- | --- | --- | --- |
| 14037654 | G | A | 40 | 17 | 1 | intergenic |  |  |
| 14193236 | G | A | 40 | 20 | 0,87 | intergenic |  |  |
| 14955241 | G | A | 40 | 20 | 0,95 | intergenic |  |  |
| 15012908 | G | A | 40 | 27 | 0,93 | intergenic |  |  |
| 15704281 | G | A | 40 | 27 | 0,84 | intergenic |  |  |
| 15708443 | G | A | 40 | 35 | 0,92 | intergenic |  |  |
| 15751944 | G | A | 40 | 31 | 0,89 | intergenic |  |  |
| 15769876 | G | A | 40 | 21 | 0,91 | intergenic |  |  |
| 15867979 | G | A | 40 | 27 | 1 | intergenic |  |  |
| 16422276 | G | A | 40 | 15 | 0,88 | intergenic |  |  |
| 16610140 | G | A | 40 | 27 | 1 | intergenic |  |  |
| 16857756 | G | A | 40 | 19 | 1 | intergenic |  |  |
| **17308283** | **G** | **A** | **40** | **25** | **0,89** | **splice site** | **AT1G47230** | **CYCLIN A3;4 (CYCA3;4)** |
| 17314228 | G | A | 40 | 16 | 1 | intergenic |  |  |
| 17346438 | G | A | 40 | 18 | 1 | intergenic |  |  |
| 17793250 | G | A | 40 | 20 | 0,95 | intronic noncoding | AT1G48180 | target of trans acting-siR480/255 protein |
| 17923246 | G | A | 40 | 17 | 0,89 | intronic noncoding | AT1G48490 | INCOMPLETE ROOT HAIR ELONGATION 3 (IRE3) |

**Supplemental Table 2.**
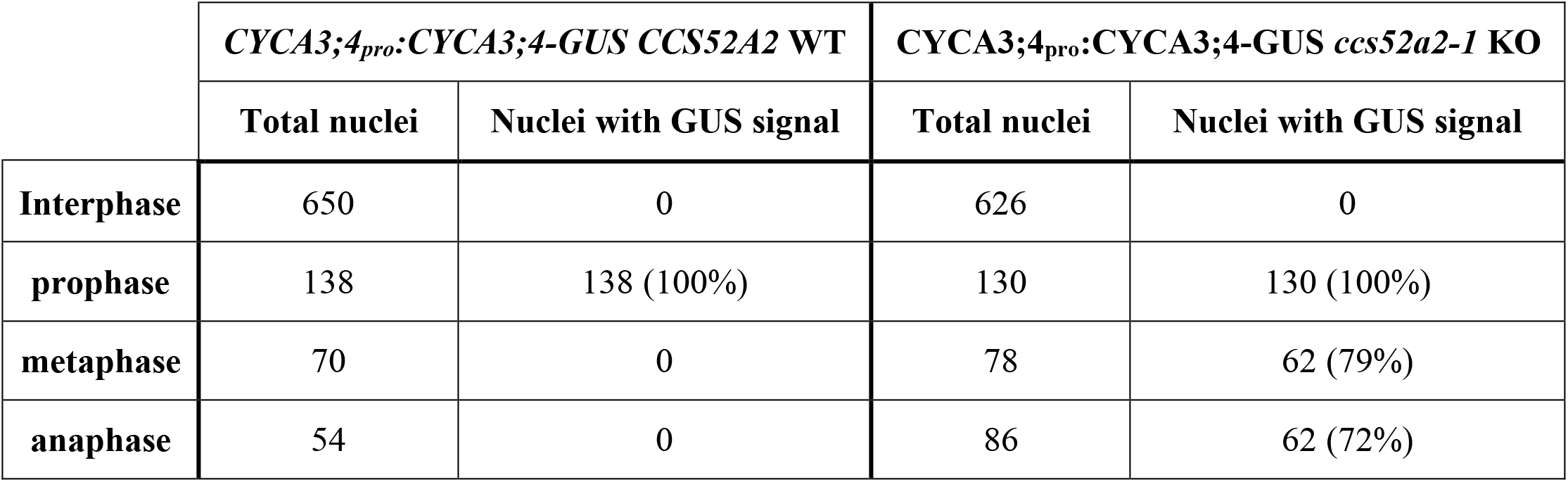
Number of Nuclei Showing CYCA3;4-GUS Signal Throughout the Cell Cycle in Root Tips With and Without Functional *CCS52A2*.

|  | *CYCA3;4<sub>pro</sub>:CYCA3;4-GUS CCS52A2* WT |  | *CYCA3;4<sub>pro</sub>:CYCA3;4-GUS ccs52a2-1* KO |  |
| --- | --- | --- | --- | --- |
|  | Total nuclei | Nuclei with GUS signal | Total nuclei | Nuclei with GUS signal |
| **Interphase** | 650 | 0 | 626 | 0 |
| **prophase** | 138 | 138 (100%) | 130 | 130 (100%) |
| **metaphase** | 70 | 0 | 78 | 62 (79%) |
| **anaphase** | 54 | 0 | 86 | 62 (72%) |

**Supplemental Table 3.**
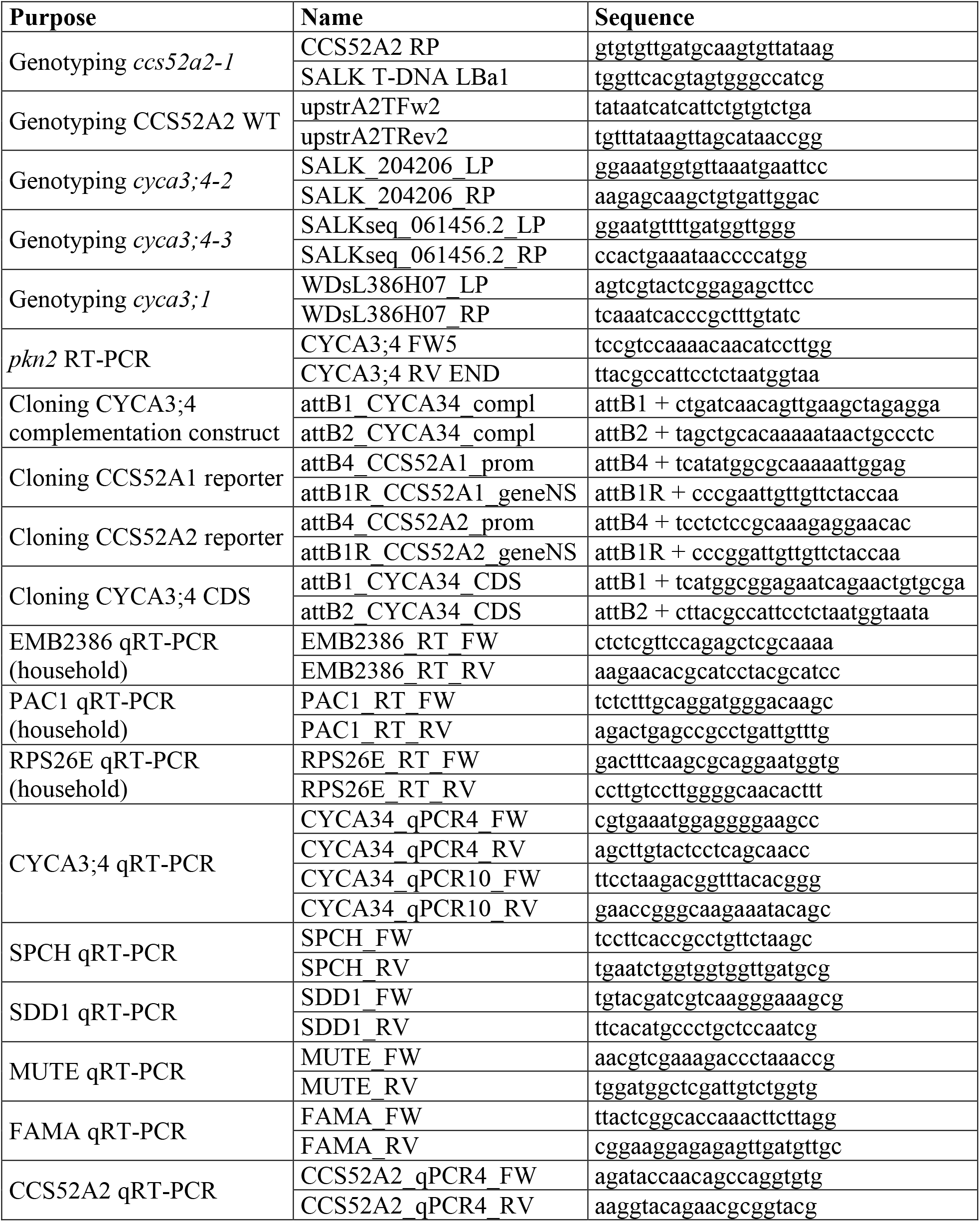
Primer Sequences

